# Centrocortin potentiates co-translational localization of its mRNA to the centrosome via dynein

**DOI:** 10.1101/2024.08.09.607365

**Authors:** Hala Zein-Sabatto, Jovan S. Brockett, Li Jin, Christian A. Husbands, Jina Lee, Junnan Fang, Joseph Buehler, Simon L. Bullock, Dorothy A. Lerit

**Author notes:** University of Pennsylvania School of Dental Medicine, Philadelphia, PA 19104. co-corresponding authors **Correspondence:** Simon L. Bullock, Ph.D. Dorothy A. Lerit, Ph.D. **Lead contact:** Dorothy A. Lerit, Ph.D.

## Abstract

Centrosomes rely upon proteins within the pericentriolar material to nucleate and organize microtubules. Several mRNAs also reside at centrosomes, although less is known about how and why they accumulate there. We previously showed that local *Centrocortin* (*Cen*) mRNA supports centrosome separation, microtubule organization, and viability in *Drosophila* embryos. Here, using *Cen* mRNA as a model, we examine mechanisms of centrosomal mRNA localization. We find that while the Cen N’-terminus is sufficient for protein enrichment at centrosomes, multiple domains cooperate to concentrate *Cen* mRNA at this location. We further identify an N’-terminal motif within Cen that is conserved among dynein cargo adaptor proteins and test its contribution to RNA localization. Our results support a model whereby Cen protein enables the accumulation of its own mRNA to centrosomes through a mechanism requiring active translation, microtubules, and the dynein motor complex. Taken together, our data uncover the basis of translation-dependent localization of a centrosomal RNA required for mitotic integrity.

**Summary:** Enrichment of *Centrocortin* (*Cen*) mRNA at centrosomes is required for mitotic fidelity. This study describes a mechanism underlying co-translational *Cen* mRNA targeting involving microtubules, the dynein motor, and a highly conserved dynein binding motif within the *Cen* coding sequence.

## Introduction

RNA localization is a highly conserved paradigm used to restrict gene expression to subcellular compartments (Chin and Lecuyer, 2017; Ryder and Lerit, 2018; Das et al., 2021). Several mechanisms enable RNA localization, including active transport, selective protection from degradation, and diffusion coupled to local entrapment (Palacios, 2007; Holt and Bullock, 2009; Das et al., 2021). Based on a small number of well-characterized examples, such as *β-actin* mRNA, it is widely believed that active transport involves recognition of RNA elements by RNA-binding proteins, which then recruit motor proteins to traffic the mRNA cargo to its destination (Kislauskis et al., 1993; Oleynikov and Singer, 2003; Bullock, 2007; Martin and Ephrussi, 2009; Mofatteh and Bullock, 2017). Often, these RNAs are translated once they reach their destination (Besse and Ephrussi, 2008; Jung et al., 2014). However, in cases of co-translational transport, the nascent peptide plays a critical role in RNA localization, as classically shown for transcripts localizing to the endoplasmic reticulum (recently reviewed in (Gasparski et al., 2022)).

Recent work highlights the centrosome as a subcellular hub for mRNA localization and translational control (Marshall and Rosenbaum, 2000; Lecuyer et al., 2007; Ryder and Lerit, 2018; Zein-Sabatto and Lerit, 2021). Centrosomes undergo cell cycle-dependent oscillations in microtubule-organizing activity dependent upon the recruitment and shedding of the pericentriolar material (PCM) (Gould and Borisy, 1977; Khodjakov and Rieder, 1999; Palazzo et al., 2000). Whether local RNAs contribute to centrosome dynamics or function is a longstanding question subject to renewed interest (Zein-Sabatto and Lerit, 2021; Lerit, 2022).

Localization-based screens in cultured cells, *Xenopus*, *Drosophila*, and other systems identified several conserved mRNAs residing at centrosomes, including *cyclin B* (*cyc B*), *Pericentrin* (*pcnt*)/ *Pcnt-like protein* (*Plp*), and *Centrocortin* (*Cen*) mRNAs (Raff et al., 1990; Groisman et al., 2000; Lecuyer et al., 2007; Sepulveda et al., 2018; Bergalet et al., 2020; Chouaib et al., 2020; Ryder et al., 2020; Safieddine et al., 2021; Fang and Lerit, 2022). Intriguingly, most RNAs enrich at centrosomes just prior to mitotic onset, with lower levels detected during M-phase (Sepulveda et al., 2018; Ryder et al., 2020). These findings suggest the concentration of RNA at the centrosome is dynamically regulated, perhaps through conserved mechanisms, and further hint at biological relevance.

Within syncytial *Drosophila* embryos, RNA localization to centrosomes is also regulated developmentally. *Drosophila* embryos proceed through 14 abridged and synchronous nuclear divisions prior to cellularization (Foe and Alberts, 1983). Most localized RNAs progressively enrich at interphase centrosomes as the nuclear cycles (NCs) proceed (Ryder et al., 2020; Fang and Lerit, 2022). For example, during NC 10, *Cen* mRNA localizes to centrosomes primarily as single molecules. However, by NC 13, significantly more *Cen* mRNA enriches at centrosomes within distinct, micron-scale ribonucleoprotein (RNP) granules containing *Cen* mRNA and protein and the multifunctional RNA-binding protein, fragile-X mental retardation protein (FMRP), a negative regulator of *Cen* mRNA translation (Ryder et al., 2020).

Cen was originally identified based on its direct binding to Centrosomin (Cnn), an essential PCM scaffolding factor (Kao and Megraw, 2009). That study further showed that *Cen* mutants display mitotic errors and embryonic lethality. Critically, proper localization of *Cen* mRNA to the centrosome is also important for mitotic fidelity. The 3’-UTR of the anterior morphogen *bicoid* (*bcd*) contains localization elements sufficient to direct heterologous RNAs to the anterior pole (Macdonald and Struhl, 1988). By fusing the *Cen* coding sequence (CDS) to the *bcd* 3’-UTR, we demonstrated that *Cen* mRNA mislocalization results in centrosome separation errors, disorganized microtubules, DNA damage, and embryonic lethality (Ryder et al., 2020). What directs *Cen* mRNA to the centrosome remains little understood, however.

Because the early *Drosophila* embryo is largely transcriptionally quiescent, and its development relies upon maternally endowed stores of RNAs and proteins until the maternal-to-zygotic transition (Tadros and Lipshitz, 2009), the rapid accumulation of RNA at interphase centrosomes is suggestive of an active transport mechanism. However, this remains to be tested. We and others showed the *Cen* CDS is necessary and sufficient for RNA localization (Bergalet et al., 2020; Ryder et al., 2020). Consistent with a targeting mechanism requiring the nascent peptide, the accumulation of *Cen* mRNA at centrosomes is sensitive to the protein synthesis inhibitor harringtonine (Bergalet et al., 2020). This finding also aligns with the discovery that mRNAs localize to centrosomes in mammalian cells while they are translated (Safieddine et al., 2021). While co-translational transport has emerged as the prevailing model for all centrosome-localized mRNAs studied to date, the underlying mechanisms directing these RNAs to the centrosome remains largely unknown.

Here, we investigate how *Cen* RNPs localize to the centrosome. We identify *cis-* and *trans*-elements needed for proper localization of *Cen* mRNA. We show that an N-terminal Cen fragment is sufficient for protein localization to the centrosome but insufficient to form and localize *Cen* mRNA granules. Nevertheless, the N’-terminus of Cen is necessary for the accumulation of *Cen* mRNA at centrosomes. Our data indicate that multiple domains within the *Cen* CDS work together to coordinate effective RNA localization. Supporting this notion, we identified a conserved dynein light intermediate chain (DLIC) binding site within the Cen N-terminus that, together with other components of the dynein motor complex, promotes *Cen* mRNA granule organization and localization. We propose a model whereby Cen protein serves as a dynein cargo adaptor to potentiate the co-translational localization of its cognate mRNA.

## Results

### Engaged polysomes are necessary for *Cen* mRNA granule formation and localization to centrosomes

*Cen* mRNA localization displays differential sensitivity to various classes of translational inhibitors (Bergalet et al., 2020). Using single molecule fluorescence *in situ* hybridization (smFISH) and computational analysis of the resulting images (Ryder et al., 2020; Ryder and Lerit, 2020), we quantified *Cen* mRNA localization relative to GFP-γTubulin (GFP-γTub) labelled centrosomes in embryos treated with puromycin (puro), a tRNA analog that terminates translation elongation and promotes ribosome dissociation, versus anisomycin (aniso) and cycloheximide (CHX), drugs that block elongation without releasing the nascent peptide (Figure 1A) (Nathans, 1964; Grollman, 1967; Schneider-Poetsch et al., 2010). Each of the translational inhibitors we examined impaired *Cen* mRNA accumulation at centrosomes, also resulting in a corresponding reduction in the percent of RNA localizing within higher order RNP granules (defined as four or more overlapping RNA objects (Ryder et al., 2020)) (Figure 1B,C). These responses were particularly evident upon treatment with puro, where RNA localization and granule formation were largely abolished. Thus, *Cen* mRNA localization is dependent upon intact polysomes. Our findings further suggest that sequences within the nascent peptide direct *Cen* mRNA localization.

**Figure 1.**
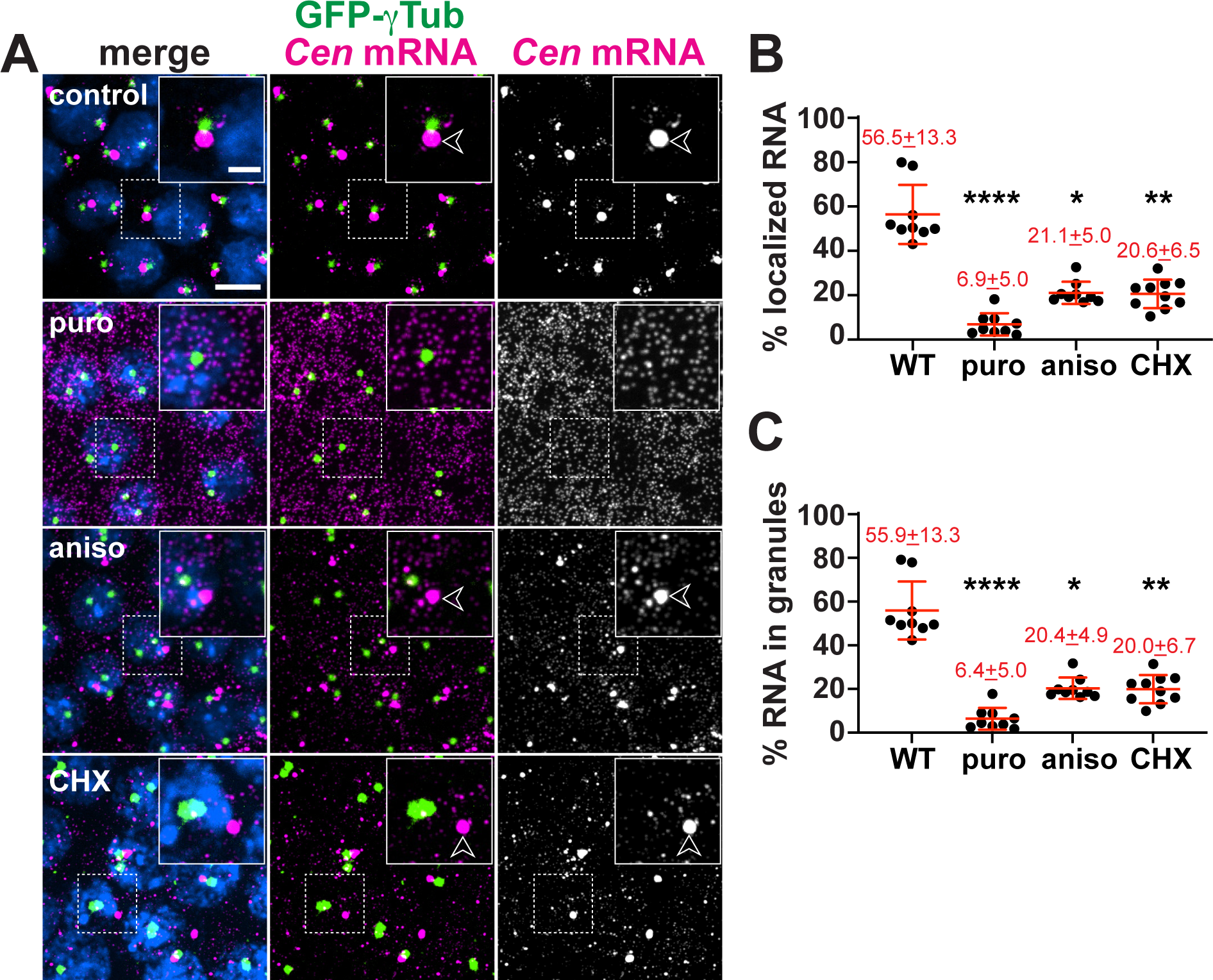
Co-translational transport of *Cen* mRNA to centrosomes. (A) Maximum-intensity projections of NC 13 embryos expressing GFP-γTub (green) stained with *Cen* smFISH probes (magenta) and DAPI (blue) to label nuclei following incubation with DMSO (control) or the translation inhibitors puromycin (puro), anisomycin (aniso), or cycloheximide (CHX). Arrowheads mark *Cen* RNPs. Quantification shows the percentage of *Cen* mRNA (B) localizing to the centrosome and (C) organized within granules, defined as ≥4 overlapping RNA objects (Ryder et al., 2020). Mean ± SD is displayed (red). Significance by ANOVA with Dunnett’s multiple comparison test with *, P<0.05; **, P<0.01; and ****, P<0.0001. Scale bars: 5 µm; 2 µm (insets).

### The Cen N-terminus is necessary, but not sufficient, to localize *Cen* RNA to centrosomes

To identify domains important for *Cen* mRNA localization, we first truncated the Cen protein into N-(*CenΔC*, comprising amino acids (AAs) 1–289) or C-terminal (*CenΔN*, comprising AAs 290– 790) pieces and expressed these in the *Cen* null genetic background (Figure 2A).

**Figure 2.**
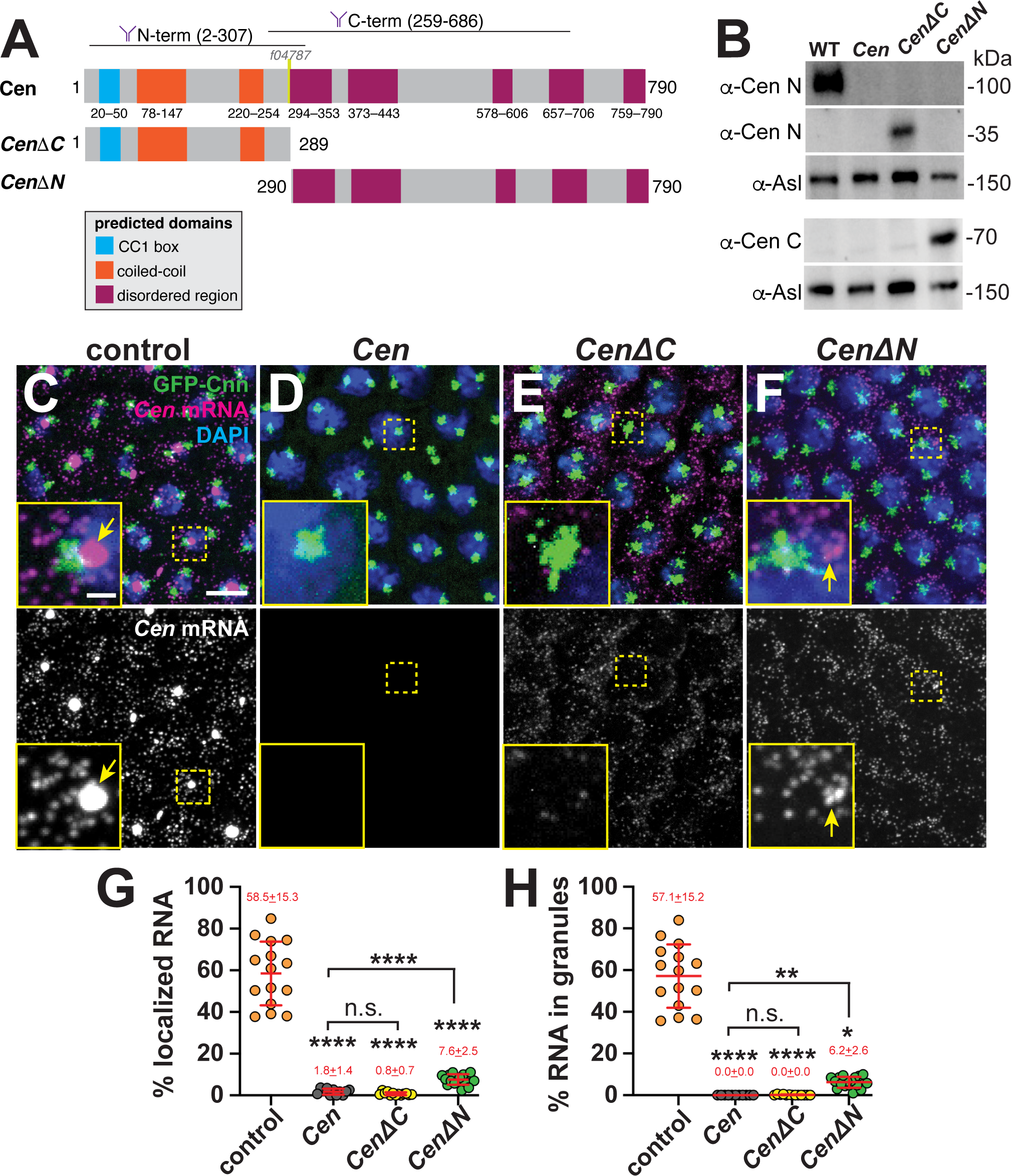
Multiple Cen domains support mRNA localization to the centrosome. (A) Schematic of the full-length and truncated Cen protein products with positions of predicted domains (Paysan-Lafosse et al., 2023), antibody epitopes (Kao and Megraw, 2009), and the transposon *f04787* within null mutants indicated. **(B)** Immunoblots from 0.5–2.5 hr embryo extracts from the indicated genotypes showing truncated Cen protein products in the *CenΔC* (∼35 kDa) and *CenΔN* (∼70 kDa) samples relative to the Asl loading control. The N-terminal anti-Cen antibody was used for the top two blots (α-Cen N), while the C-terminal anti-Cen antibody was used below (α-Cen C; see also (Kao and Megraw, 2009)). (**C–F**) Maximum-intensity projections of *Cen* smFISH (magenta) in NC 13 interphase embryos expressing *GFP-Cnn* (green) with DAPI-stained nuclei (blue). **(C)** Control embryo with *Cen* mRNA localized at centrosomes (arrow). In contrast, **(D)** *Cen* mutants and (**E**) *CenΔC* embryos fail to localize *Cen* mRNA to centrosomes. (**F**) Although *CenΔN* is partially sufficient to form small RNA granules (arrow) near centrosomes, neither fragment recapitulates WT localization. In all experiments, *CenΔC* and *CenΔN* are expressed in the *Cen* null background. Percentage of *Cen* mRNA **(G)** overlapping with centrosomes or **(H)** in granules 0 µm from the Cnn surface. Each dot represents a measurement from N= 15 control, 11 *Cen*, 13 *CenΔC*, and 17 *CenΔN* embryos. Mean ± SD is displayed (red). Significance was determined by (G) one-way ANOVA followed by Dunnett’s T3 multiple comparison test or (H) Kruskal-Wallis test followed by Dunn’s multiple comparison test with n.s., not significant; *, P<0.05; **, P<0.01; and ****, P<0.0001. Scale bars: 5µm; 1µm (insets).

Immunoblotting confirmed the truncated products were expressed at comparable levels in early embryo extracts and migrated at the expected molecular sizes, as detected by antibodies with epitopes in the N’-or C’-regions of Cen (Figure 2B; (Kao and Megraw, 2009)). By interphase of NC 13, most *Cen* mRNA normally localizes to the centrosome within granules (*arrows*, Figure 2C). Demonstrating specificity, the smFISH signals were absent in *Cen* mutants, which harbor a P-element insertion (*f04787*) in the CDS and are RNA and protein nulls (Figure 2A,D; (Bergalet et al., 2020; Ryder et al., 2020)). Because our probes tile the CDS, *Cen* smFISH signals were detected in both *CenΔC* and *CenΔN* backgrounds (Figure 2E and F). However, the percentages of *Cen* mRNA overlapping the centrosome surface (0 µm distance from GFP-Centrosomin (GFP-Cnn)) and within granules were dramatically reduced in the truncation lines relative to controls (Figure 2G and H). While small RNA granules were occasionally detected in *CenΔN* embryos (*arrows*, Figure 2F, H), neither fragment was sufficient to restore endogenous levels of RNA localization. We conclude that neither the N’-nor C’-termini of Cen are sufficient for robust RNA localization; rather, both regions likely function cooperatively.

Cen protein also localizes to the centrosome within *Cen* RNPs (Bergalet et al., 2020; Ryder et al., 2020). Therefore, we next tested whether the truncated protein products localized to centrosomes. We confirmed that anti-Cen N’ and anti-Cen C’ antibodies detect Cen at centrosomes in control embryos, and these signals are absent in *Cen* null mutants (Figures 2A and 3A–C, F; (Kao and Megraw, 2009)). By comparison, while CenΔC localized to centrosomes, CenΔN did not, as detected by the N’-versus C’-terminal antibodies, respectively (Figure 3D– F). Contrary to full-length Cen, we observed that the CenΔC protein appeared to localize near the center of the centrosome rather than the outer PCM flares (*cf.* Figure 3A, B vs. D); however, what directs Cen to distinct PCM zones remains unclear. We conclude that the N-terminus is necessary and sufficient for Cen protein localization (Figure 3D–F). Moreover, our results show that *Cen* mRNA and protein distributions may be uncoupled (*cf.* Figures 2E vs. 3D).

**Figure 3.**
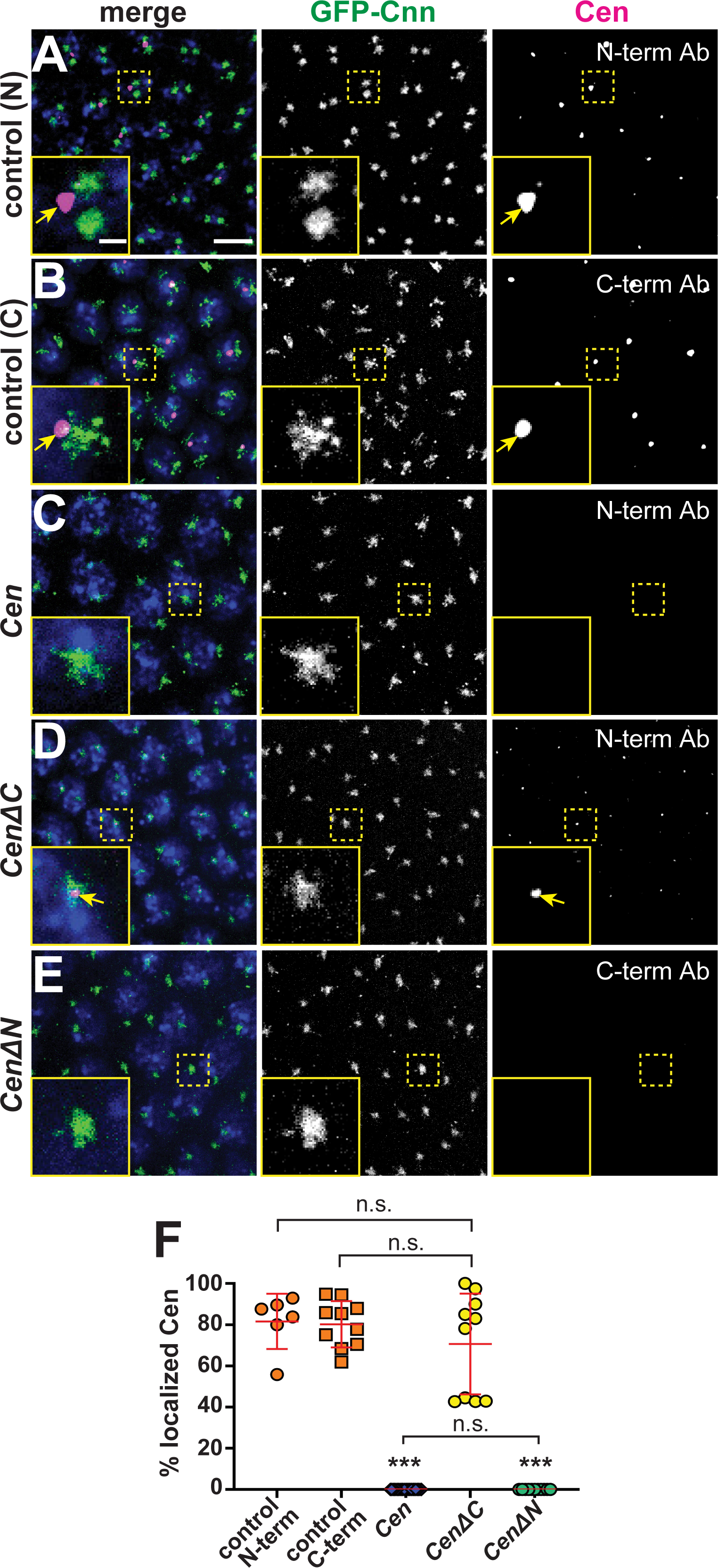
The N-terminal fragment is necessary and sufficient for Cen protein localization to the centrosome. Maximum-intensity projections of NC 13 interphase embryos expressing *GFP-Cnn* (green) labeled with anti-Cen antibodies (magenta) and DAPI (blue nuclei). Control embryos labeled with **(A)** anti-Cen N-terminal or (**B**) C-terminal antibodies (Ab) show Cen localized at centrosomes (arrows). **(C)** *Cen* protein is not detected in null mutants. **(D)** The N-terminal fragment (*CenΔC*) is sufficient to direct Cen to the centrosome (arrows), while the C-terminal fragment (*CenΔN*; (**E**)) is not. Both transgenes are expressed in the *Cen* null background. **(F)** The percentage of Cen protein signals overlapping with centrosomes (0 µm from Cnn surface). Each dot represents a measurement from N= 6 control (N-terminal Cen Ab), 10 control (C-terminal Cen Ab), 23 *Cen* null (N-terminal Cen Ab), 10 *CenΔC* (N-terminal Cen Ab), and 11 *CenΔN* embryos (C-terminal Cen Ab). Significance was determined by Kruskal-Wallis test followed by Dunn’s multiple comparison test with n.s., not significant and ***, P<0.001. Scale bars: 5µm; 1µm (insets).

As an independent approach to experimentally uncouple *Cen* mRNA and protein, we deleted the translation initiation codon (*Cen^-ATG^*) and expressed this or a full-length control (*Cen^FL^*) HA-tagged transgene in the *Cen* null background. Unexpectedly, *Cen^-ATG^*was translated in ovaries and embryos, yielding a protein ∼30 kDa smaller than *Cen^FL^*, as detected by western blotting (Figure 4A). These data indicate Cen has one or more cryptic translation start sites. Consistent with this finding, the first 90–100 N’-terminal AAs of Cen^-ATG^ were undetectable by mass spectrometry (Figures S1 and 4B). Although *Cen^-ATG^* did not block Cen translation as intended, it did permit further analysis of the role of the N’-terminus for *Cen* mRNA localization. The amount of *Cen* mRNA localizing to centrosomes was reduced by two-thirds and granules failed to form in *Cen^-ATG^* embryos relative to *Cen^FL^* controls (Figure 4C–F). In addition, despite comparable expression levels, less Cen^-ATG^ protein localized to centrosomes compared to Cen^FL^ (*insets* Figure 4C, D). Taken together, our analysis indicates that the first ∼100 AA are important for *Cen* mRNA and protein localization to centrosomes.

**Figure 4.**
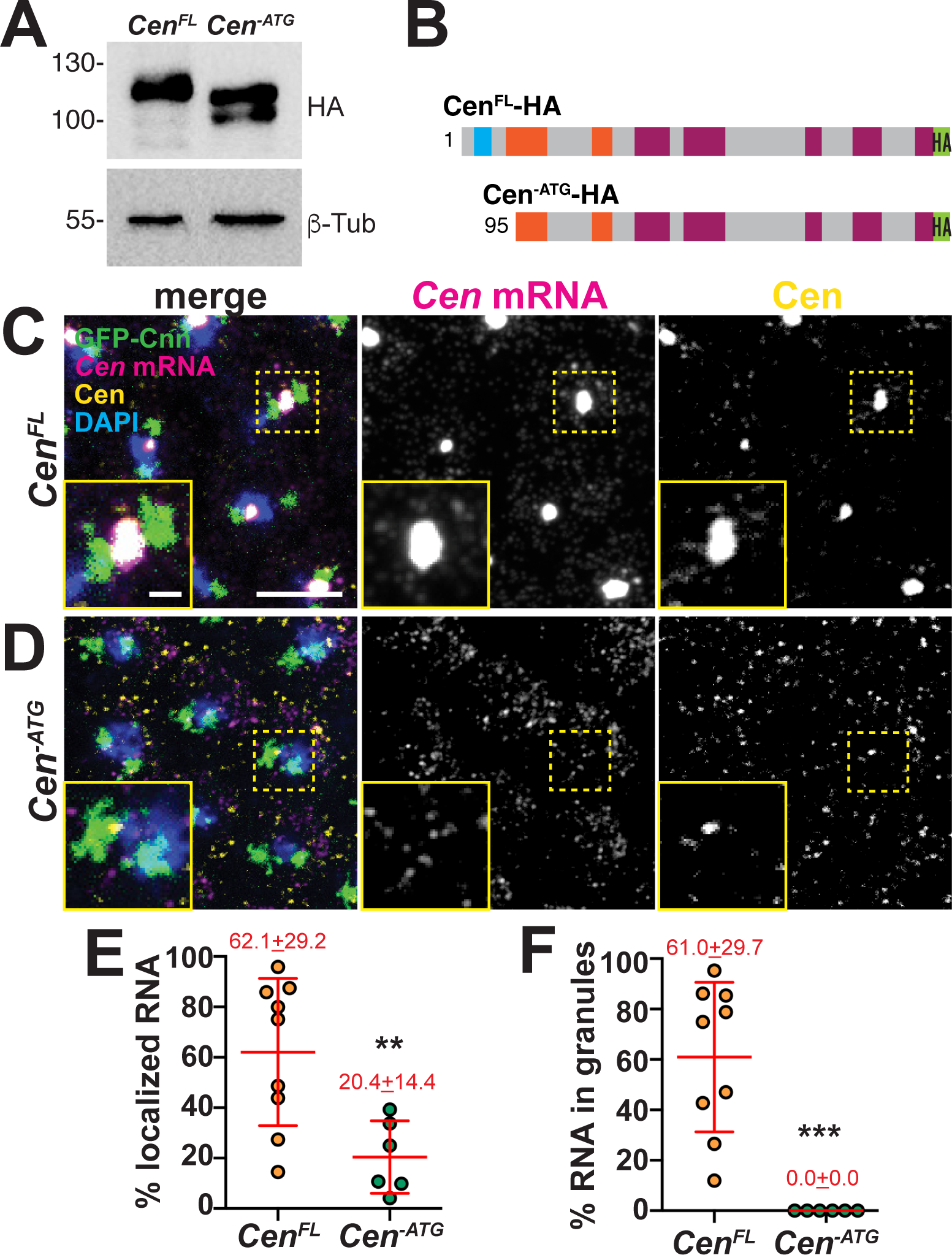
The first 100 AA of Cen direct RNA localization. (**A**) Immunoblots from ovarian extracts from the indicated genotypes showing Cen protein products, as detected with anti-HA antibodies, relative to the B-Tub loading control. Truncated products are detected in the *Cen^-ATG^* lysate. (**B**) Schematic of the *Cen^FL^* and *Cen^-ATG^* HA-tagged protein products showing predicted translation start sites, based on mass spectrometry analysis (*see Figure S1*). Maximum intensity projections of NC 13 (**C**) *Cen^FL^* and (**D**) *Cen^-ATG^* embryos expressing *GFP-Cnn* and stained with *Cen* smFISH probes (magenta), C-term anti-Cen antibodies (yellow), and DAPI (blue) to label nuclei. Quantifications show (**E**) the percentage of RNA overlapping with centrosomes or (**F**) organized within granules 0 µm from the Cnn surface. Each dot represents a measurement from N= 9 *Cen^FL^* and 6 *Cen^-ATG^* embryos. In all experiments, both transgenes were expressed in the *Cen* null background. Mean ± SD is displayed (red). Significance was determined by two-tailed Mann-Whitney test with **, p=0.0076 and ***, p=0.0004. Scale bars: 5µm; 1µm (insets).

### A conserved predicted DLIC binding motif facilitates *Cen* mRNA localization to centrosomes

Analysis of the Cen protein secondary structure revealed two predicted N’-terminal coiled-coil domains and several disordered regions clustered at the C-terminus (Cen; Figure 2A; (Apweiler et al., 2000)). Through primary sequence alignments, we identified a region within the first 50 AA of Cen that is very similar to the previously identified CC1 box motif of several dynein cargo adaptors, which include BicD family members, Spindly, and Hook proteins (*red box*, Figure 5A; (Gama et al., 2017; Lee et al., 2018)). This motif is also conserved in the human *Cen* paralogs, *CDR2L* and *CDR2* (Figure 5A). The CC1 box mediates the interaction of dynein cargo adaptors with a short helix of the DLIC subunit of the dynein motor complex. Together with adjacent coiled-coil sequences that interact with Dynein heavy chain (Dhc) and the dynein activating complex, dynactin, this interaction tethers cargo to the motor and releases dynein from its autoinhibited state (Gama et al., 2017; Lee et al., 2018; Lee et al., 2020; Chaaban and Carter, 2022). Through immunoprecipitation, we tested whether Cen also associates with DLIC in embryonic extracts. Similar to the positive control BicD, Cen co-precipitated with GFP-DLIC, but not GFP alone (Figure 5B). Taken together, these data are consistent with Cen representing a novel dynein cargo adaptor.

**Figure 5.**
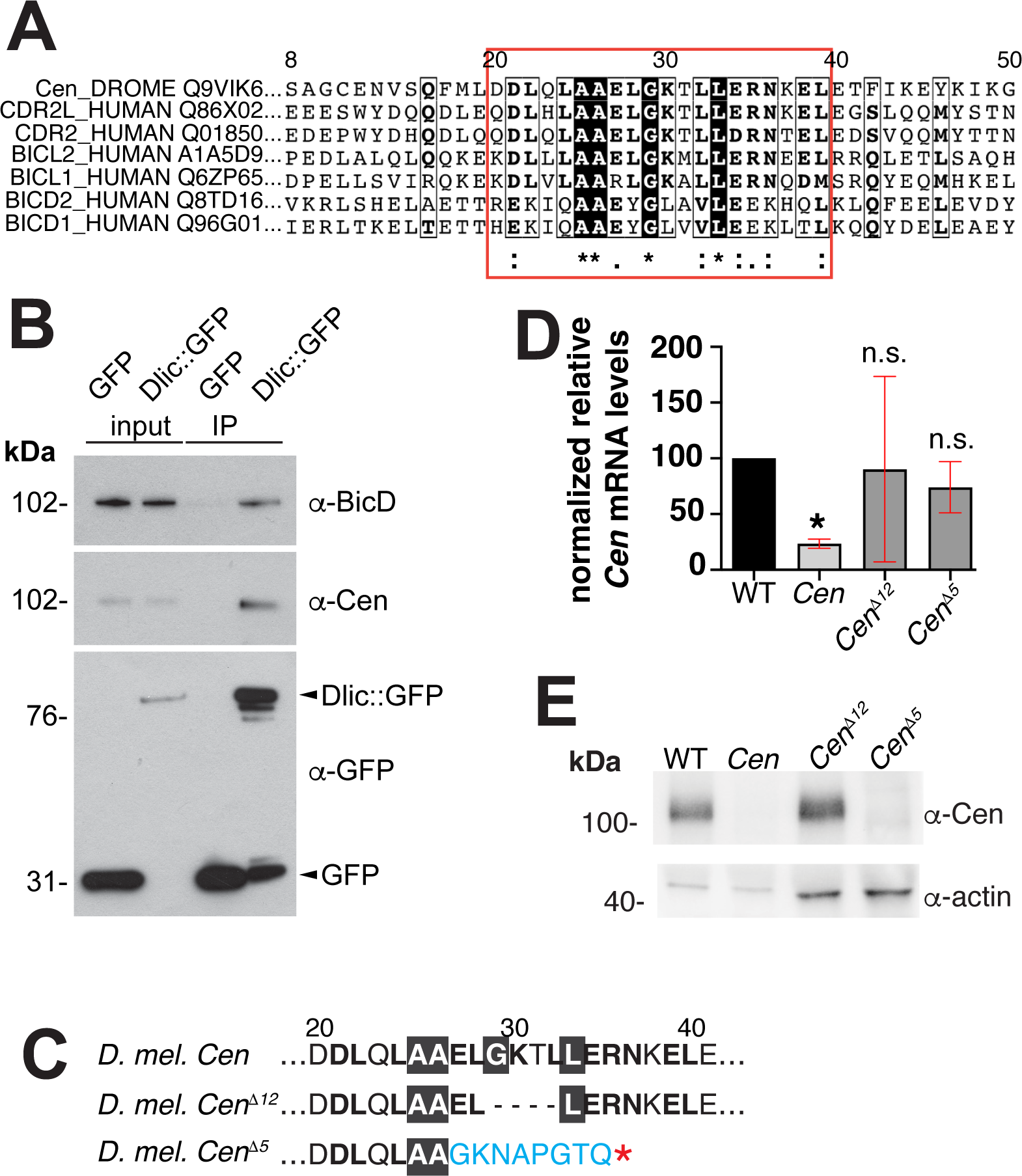
Identification of the conserved Cen DLIC binding site. (A) Clustal Omega sequence alignment of *Drosophila Cen* with the human paralogs *CDR2* and *CDR2L* and several dynein activating cargo adaptors. Red box marks the conserved DLIC-binding motif (CC1 box). (**B)** Dlic-GFP associates with BicD (Dienstbier et al., 2009) and Cen in 0–5-hour embryonic extracts. Input and immunoprecipitated samples (IP) for GFP control and Dlic-GFP are indicated. (**C**) The Cen CC1 box was mutated, yielding an in-frame deletion of the 12 nucleotides that comprise amino acids (AA) 29-32 (GKTL; *Cen^Δ12^)*, while the *Cen^Δ5^* mutant is defined by a frameshift after AA 26 and a premature stop (asterisk). **(D)** Relative levels of *Cen* mRNA normalized to *RP49* and the WT control in 0–2-hour embryos (up to NC 14) by qPCR. Bars show mean ± SD from three independent experiments. *, P<0.05 by Kruskal-Wallis multiple comparison test relative to WT; n.s., not significant. **(E)** Blot shows Cen protein detected in 0–2-hour embryos with a C’-terminal anti-Cen antibody relative to the actin loading control. No Cen protein was detected in null or *Cen^Δ5^* extracts.

To test if the conserved CC1 box supports *Cen* mRNA localization to centrosomes, we disrupted it by CRISPR/Cas-9 genome editing. We successfully generated several mutants, of which, *Cen^Δ12^* represents the largest in-frame deletion recovered and removes AAs 29-33. We also examined *Cen^Δ5^*, which causes a frameshift mutation after AA 26, resulting in a predicted truncated product (Figure 5C). Both mutations disrupt highly conserved residues with the CC1 box motif, including an invariant glycine that creates a cavity in the coiled coil for binding the DLIC helix (Lee et al., 2020). We confirmed by qPCR that both *Cen^Δ12^* and *Cen^Δ5^* mutant lines express *Cen* mRNA at levels comparable to wild-type (WT; Figure 5D). In contrast, while Cen protein is produced in *Cen^Δ12^* embryos, none was detectable in *Cen^Δ5^* extracts by western blot (Figure 5E), presumably due to protein destabilization.

To assay whether the CC1 box contributes to *Cen* mRNA localization, we compared RNA distributions in *Cen^Δ12^* and *Cen^Δ5^* embryos relative to controls. In younger, interphase NC 11 embryos, enrichments of *Cen* mRNA at centrosomes were modestly reduced in *Cen^Δ12^* embryos relative to controls, but this did not reach statistical significance. Conversely, while numerous *Cen* transcripts were still detected in *Cen^Δ5^* embryos, *Cen* mRNA localization to centrosomes was eliminated (Figure S2). These findings corroborate earlier evidence that sequences within the Cen protein are essential for *Cen* mRNA localization.

We next examined *Cen* mRNA distributions relative to the non-localizing *gapdh* mRNA during interphase NC 13, when the majority of *Cen* RNA localizes to the centrosome within granules (0 µm distance from Cnn surface; Figure 6A, D,E). Relative to controls, less *Cen* mRNA enriched at centrosomes in *Cen^Δ12^* (∼15% reduction; mean±S.D.= 60.2±12.1% in *Cen^Δ12^* versus 70.5±9.1% in controls; p=0.0343 by Kruskal-Wallis test; Figures 6A, B, and D). These data indicate that impairing the DLIC binding site compromises *Cen* mRNA localization. In contrast, the *Cen^Δ5^* mutation abolished both *Cen* mRNA localization and RNA granule assembly (Figure 6C–E). Taken as a whole, these data indicate the CC1 box motif within Cen is functionally important and further imply DLIC contributes to *Cen* mRNA localization to centrosomes.

**Figure 6.**
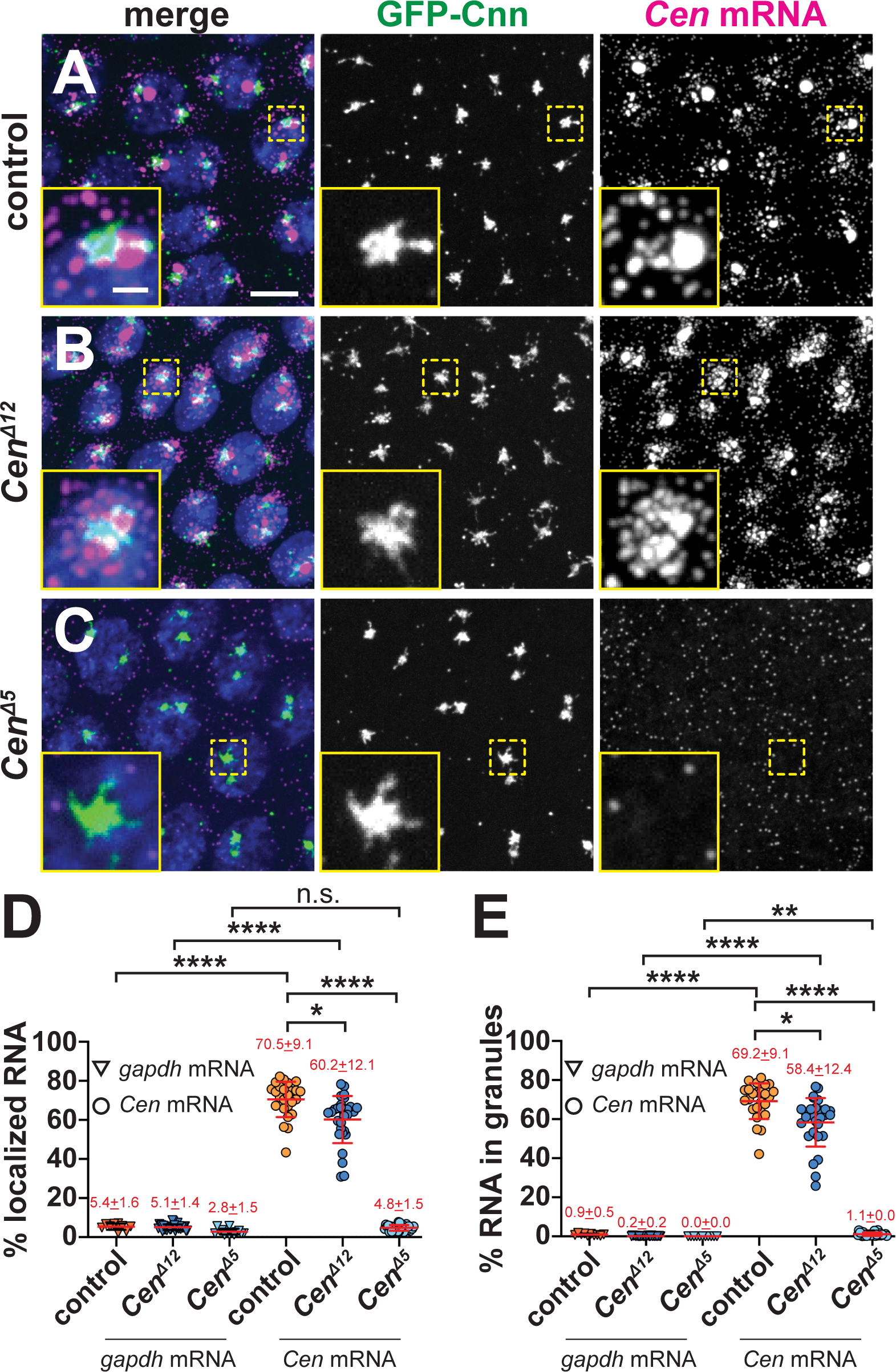
The CC1 box supports *Cen* mRNA localization. Maximum-intensity projections of NC 13 interphase embryos expressing *GFP-Cnn* (green) stained with *Cen* smFISH probes (magenta) and DAPI (blue nuclei). **(A)** Control embryos show *Cen* mRNA enriched at centrosomes in RNP granules, which are reduced in **(B)** *Cen^Δ12^* samples. **(C)** *Cen* mRNA localization and granule formation are abolished in *Cen^Δ5^* embryos. Quantification of the percentage of *Cen* or *gapdh* mRNA **(D)** overlapping with the centrosome surface and **(E)** residing in granules (0 µm distance from Cnn). Each dot represents a single measurement from control (N= 10 *gapdh* and 25 *Cen* mRNA), *Cen^Δ12^* (N= 30 *gapdh* and 30 *Cen* mRNA), and *Cen^Δ5^* (N= 14 *gapdh* and 27 *Cen* mRNA) labelled embryos. Mean ± SD displayed (red). Significance was determined by Kruskal-Wallis test followed by Dunn’s multiple comparison test relative to controls with n.s., not significant; *, P<0.05; **, P<0.01; and ****, P<0.0001. Scale bar: 5µm; 1µm (insets).

### Mitotic spindle morphogenesis is sensitive to local *Cen* mRNA dosage

To further examine the functional significance of the conserved Cen CC1 box, we examined mitotic spindle morphogenesis. Proper dosage of *Cen* mRNA at the centrosome is needed for mitotic fidelity (Ryder et al., 2020). Similar to *Cen* null mutants, *Cen^Δ12^*and *Cen^Δ5^* embryos displayed elevated frequencies of aberrant spindles and defective centrosome separation relative to controls (Figure 7A–E). These findings show that *Cen* activity supports spindle formation.

**Figure 7.**
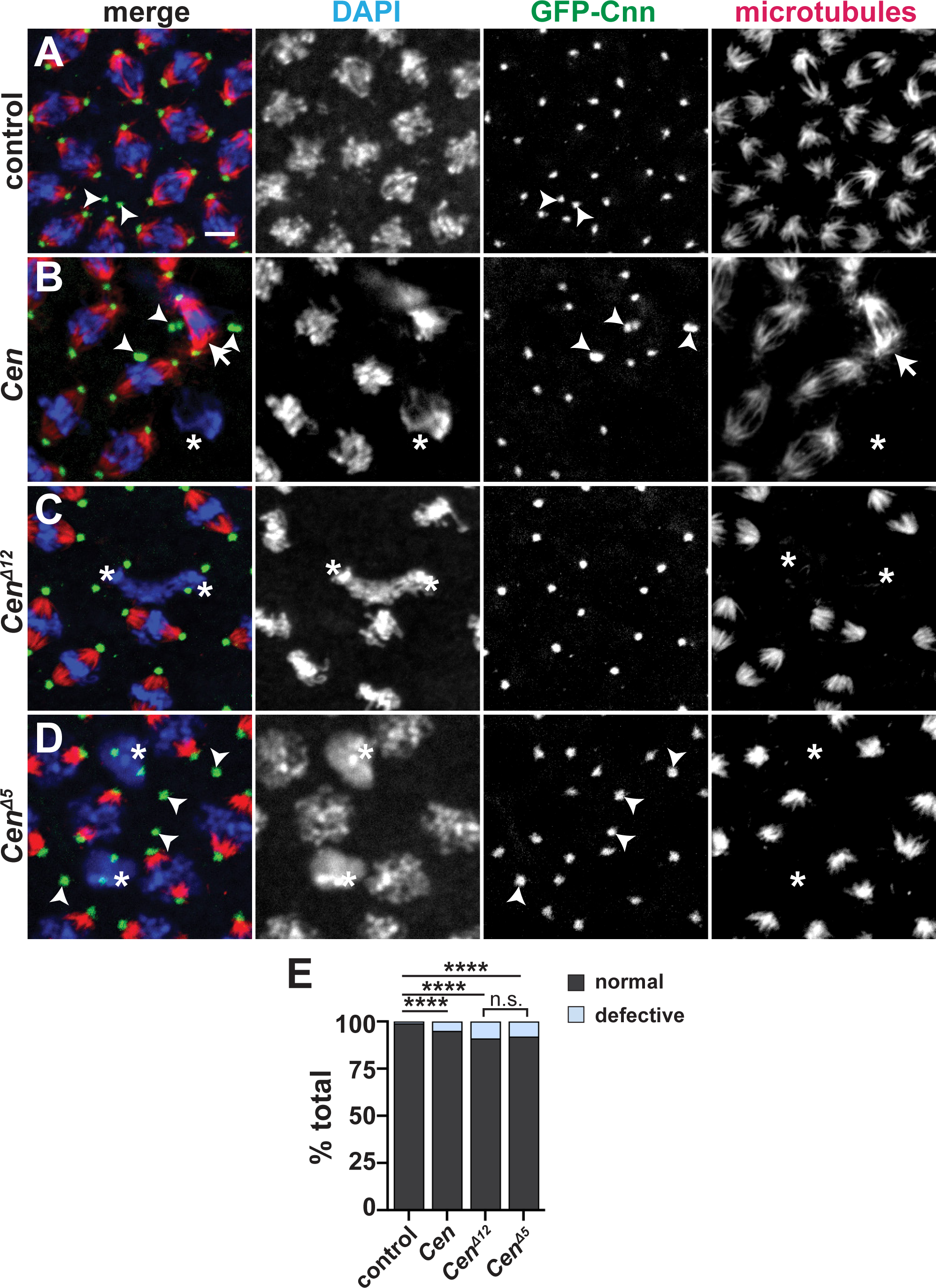
Disruption of the Cen CC1 box impairs spindle morphology. Maximum-intensity projections of metaphase NC 12 embryos from embryos expressing *GFP-Cnn* (green, centrosomes) and stained for α-Tub to label microtubules (red) and DAPI (blue nuclei). **(A)** Control embryo showing bipolar spindles. Various spindle defects are noted in **(B)** *Cen* null, (**C**) *Cen^Δ12^*, and (**D**) *Cen^Δ5^* embryos, including spindle inactivation (asterisks), detached centrosomes (arrowheads), and bent spindles (arrows). **(E)** Frequency of spindle defects from N=1622 spindles from n=7 control, N=1473 spindles from n=7 *Cen* null, n=2138 spindles from n=15 *CenΔ^12^*, and N=1842 spindles from n=12 *Cen^Δ5^* embryos. ****, P<0.00001 by Chi-square test. Scale bar: 5 µm.

### Microtubules enrich *Cen* mRNA at centrosomes

A role for the CC1 box raised the possibility that *Cen* mRNA is transported by dynein along microtubules to the centrosome. Indeed, microtubules are nucleated from centrosomes with their minus ends embedded within the PCM (Mitchison and Kirschner, 1984; Soltys and Borisy, 1985; Vertii et al., 2016). While microtubules serve as tracks for the localization of many RNAs, including *PCNT* and *ASPM* mRNAs in mammalian cells, their requirement for the localization of other centrosomal mRNAs has not been tested (Sepulveda et al., 2018; Safieddine et al., 2021).

To assay the relationship between microtubules and *Cen* mRNA, we first confirmed the coincidence of endogenous *Cen* mRNA and microtubules labeled with α-Tub antibodies by calculating a Mander’s coefficient of colocalization (Figure 8A, B; *control*). These co-occurring *Cen* mRNA and α-Tub signals were not due to spurious overlap, as rotating the RNA channel by 90° significantly reduced the extent of colocalization (Figure 8B). Therefore, a proportion of *Cen* mRNA overlaps with microtubules.

**Figure 8.**
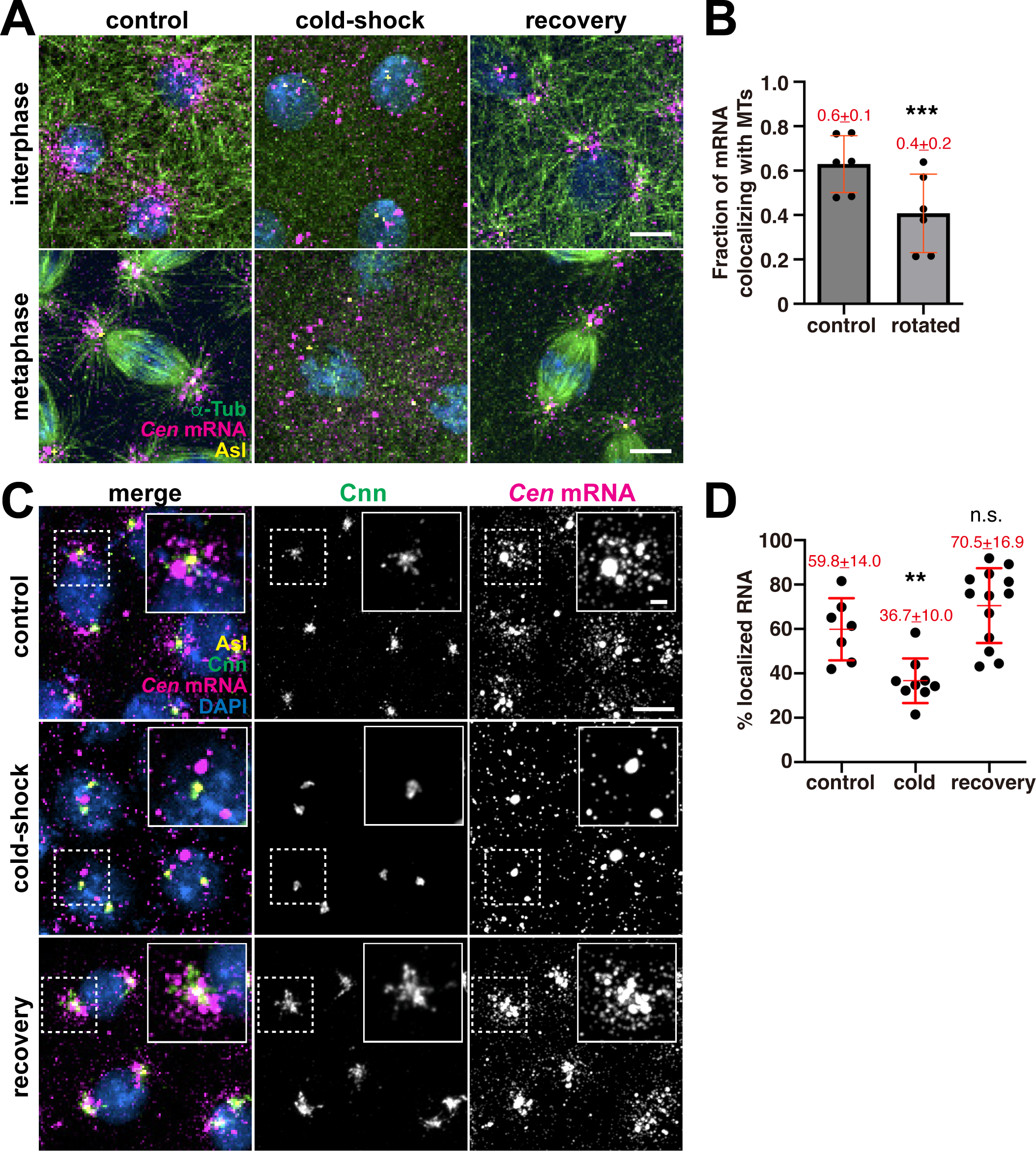
Microtubules enrich *Cen* mRNA at centrosomes. **(A)** Microtubule regrowth assay. Representative images of NC 11 embryos labeled with *Cen* smFISH probes (magenta) and antibodies for α-Tub (green) and Asl (yellow). Nuclei are labeled with DAPI (blue) in control, cold-shock, and recovery conditions. **(B)** Graph shows the Mander’s coefficient of colocalization for *Cen* mRNA overlapping with microtubules. Each dot is a measurement from N=6 interphase NC 10–11 control embryos. The RNA channel was rotated 90° to test for specificity of colocalization. **(C)** Maximum intensity projections of NC 12 interphase embryos from the indicated conditions labeled with *Cen* smFISH probes (magenta), Cnn (green) and Asl (yellow) antibodies, and DAPI (blue). Insets show Cnn structure and *Cen* mRNA distribution are affected by microtubule destabilization. **(D)** Quantification of the percentage of *Cen* mRNA localizing to centrosomes (≤1 µm distance from Asl surface) from N=7 control, 9 cold-shocked, and 13 recovered NC 12 interphase embryos. Mean ± SD is displayed (red). Significance was determined by (B) two-tailed t-test and (D) one-way ANOVA followed by Dunnett’s multiple comparisons test relative to the control with n.s., not significant; **, P<0.01; and ***, P<0.001. Scale bars: 5 µm; 2 µm (insets).

Next, we conducted a microtubule regrowth assay to determine whether microtubules are required for *Cen* mRNA localization. Cold-shock induced microtubule depolymerization was sufficient to disperse *Cen* mRNA at all stages examined (Figure 8A; *cold-shock*). Examination of RNA distributions in age-matched embryos relative to centrosomes labeled with Cnn and Asterless (Asl) antibodies revealed that microtubule disruption was sufficient to decrease *Cen* mRNA localization to centrosomes by nearly 40%, as compared to controls (Figure 8C, D). The condensation of Cnn into a more compact structure following cold-shock (Figure 8C) serves as an internal control, as identical responses were previously noted following acute microtubule depolymerization via colchicine (Megraw et al., 2002; Lerit et al., 2015). Moreover, when the embryos were allowed to briefly recover at room temperature to permit microtubule regrowth, *Cen* mRNA re-decorated microtubules and re-localized to centrosomes to untreated levels (Figure 8A; *recovery*; and C,D). These data demonstrate that microtubules allow robust localization of *Cen* mRNA to centrosomes.

### *Cen* mRNA localization is dynein-dependent

To further test whether dynein traffics *Cen* mRNA to centrosomes, we compared *Cen* distributions in embryos with impaired Dhc activity relative to controls. Because dynein is essential for viability (Gepner et al., 1996), we collected embryos from mothers homozygous for a hypomorphic mutation in the *Dhc* gene (*Dhc64C*) (Salvador-Garcia et al., 2023) that is equivalent to the *legs at odd angles* (LOA; hereafter, *Dhc^LOA^*) allele first described in mouse (Nolan et al., 2000; Hafezparast et al., 2003). Supporting a requirement of the dynein motor complex for *Cen* mRNA localization, the *Dhc^LOA^* mutation led to significantly less *Cen* mRNA at centrosomes (∼40% reduction), as well as ∼50% less *Cen* mRNA within granules, as compared to controls (Figure 9A–E). Moreover, the *Cen* RNPs that did form in the *Dhc* mutants were also smaller (Figure 9B, E’). These data confirm that dynein supports *Cen* RNA granule assembly and localization. In contrast, depletion of the plus-end-directed microtubule motor, kinesin, using a shRNA (*Khc^RNAi^*) sufficient to reduce Khc protein levels (Veeranan-Karmegam et al., 2016) did not significantly alter *Cen* RNA localization at the centrosome (Figure 9C–E). Our collective data implicate the dynein transport complex in promoting *Cen* mRNA accumulation at centrosomes.

**Figure 9.**
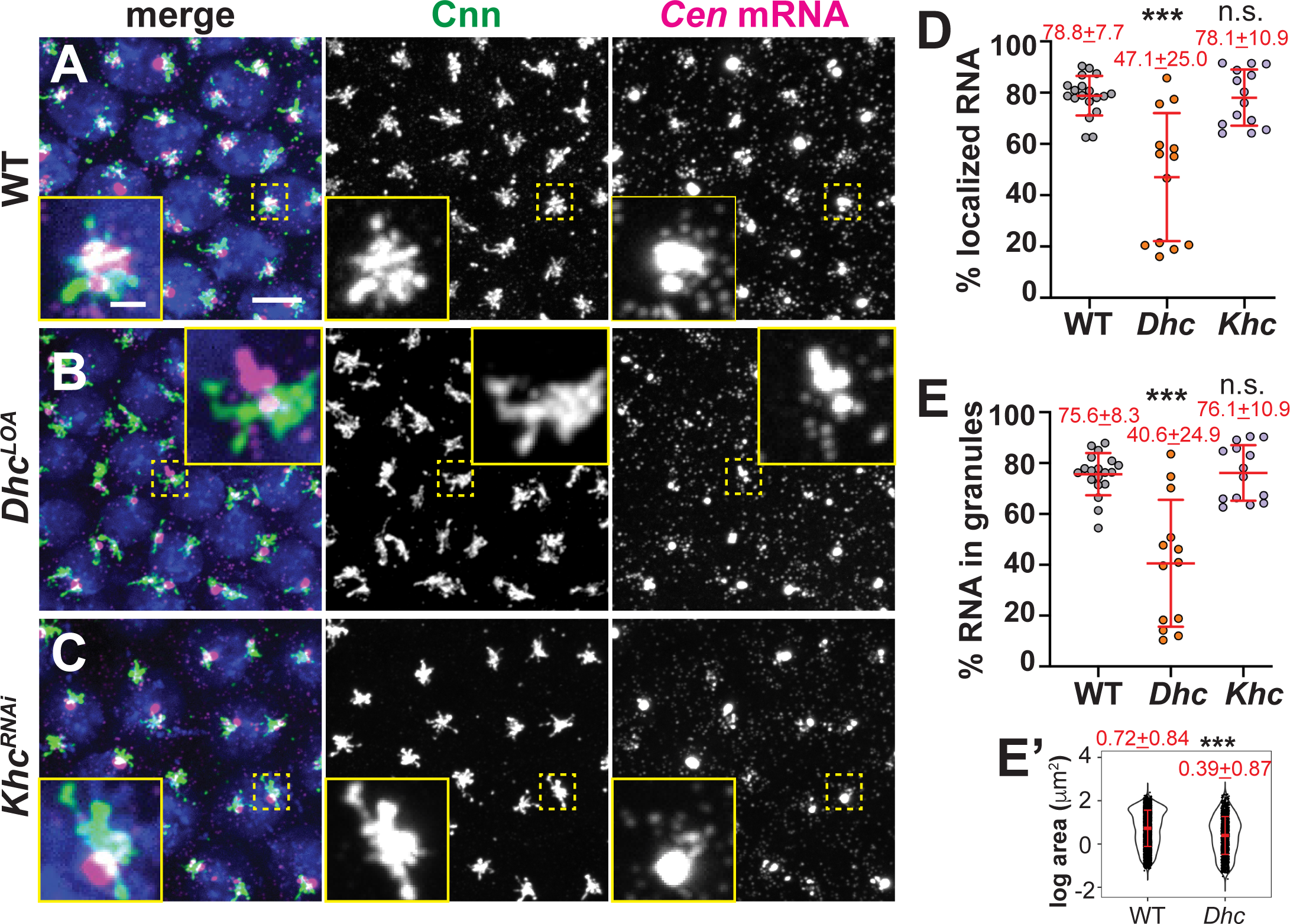
Dynein targets *Cen* mRNA to centrosomes. Maximum-intensity projections of NC 13 interphase embryos labeled with *Cen* smFISH (magenta), anti-Cnn antibodies (green; centrosomes), and DAPI (blue nuclei) in **(A)** WT, **(B)** *Dhc^LOA^* hypomorphic, or **(C)** *Khc^RNAi^* embryos. Quantification shows the percentage of total mRNA that **(D)** overlaps with centrosomes and **(E)** resides in granules at centrosomes (0 µm distance from Cnn). Each dot represents a measurement from N= 19 WT, 13 *Dhc^LOA^* and 14 *Khc^RNAi^* embryos. **(E’)** Log transformed RNA granule area from N=4127 granules from n=23 WT embryos and N=1412 granules from n=13 *Dhc^LOA^* embryos; each dot represents a single granule. Mean ± SD displayed (red). Significance by (D and E) Kruskal-Wallis test followed by Dunn’s multiple comparison test relative to WT and (E’) unpaired t-test with n.s., not significant and ***, P<0.001. Scale bar: 5µm; 1 µm (inset).

### The RNA-binding protein Egl enhances *Cen* mRNA localization

To direct RNA trafficking along microtubules in *Drosophila* oocytes and blastoderm embryos, the RNA-binding protein Egl loads various transcripts onto dynein (Dienstbier et al., 2009). Dynein light chain binds Egl and promotes its dimerization, which optimizes Egl binding to mRNA and subsequently the dynein cargo adaptor BicD (McClintock et al., 2018; Sladewski et al., 2018; Goldman et al., 2019). Egl is required for oocyte specification and polarization, precluding analysis of *egl* null embryos (Mach and Lehmann, 1997; Navarro et al., 2004). Therefore, to test whether Egl contributes to *Cen* mRNA localization, we examined *egl*-deficient embryos expressing an *egl* mutant transgene (*Egl^RBD3^*) that contains alanine substitutions of 8 positively charged residues within the Egl RNA-binding domain (RBD) and, consequently, disrupts subcellular localization of various mRNA cargoes in *Drosophila* oocytes (Goldman et al., 2021). As a control, we also examined *egl*-deficient embryos expressing a full-length (*Egl^WT^)* transgene. This analysis revealed that interphase NC 13 *Egl^RBD3^*embryos had ∼15% less *Cen* mRNA at the centrosome or within RNA granules than *Egl^WT^* (Figure 10A–D). Additionally, those *Cen* RNPs that formed in the absence of Egl RNA-binding activity were nearly 20% smaller than controls (Figure 10D’). Thus, Egl contributes to *Cen* mRNA granule assembly or maintenance to promote the accumulation of *Cen* mRNA at centrosomes.

**Figure 10.**
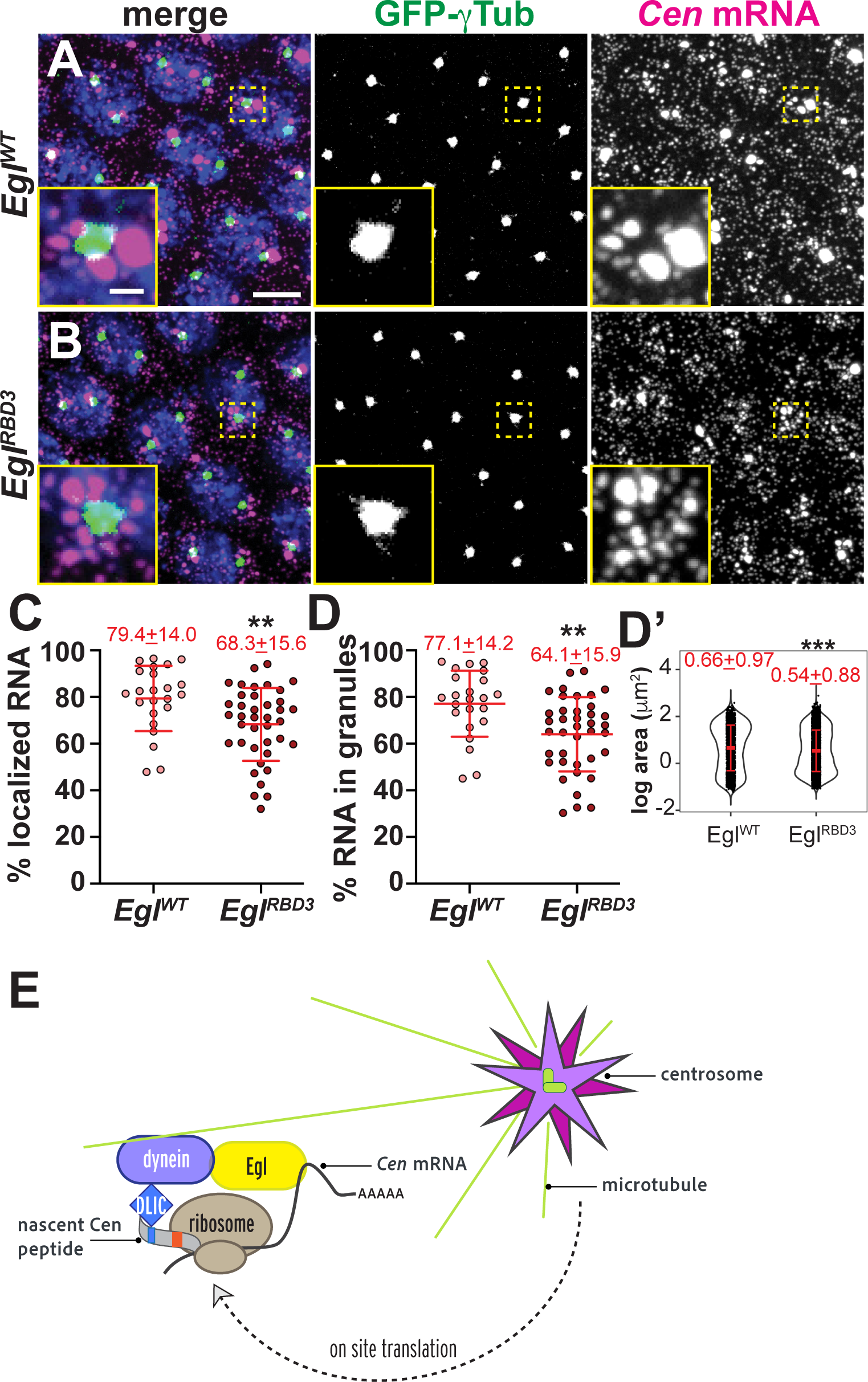
The Egl RBD supports *Cen* mRNA localization. Maximum-intensity projections of NC 13 interphase embryos expressing *GFP-γ-Tub* (green) labeled with *Cen* smFISH (magenta) and DAPI (blue) in **(A)** control *Egl^WT^* and **(B)** *Egl^RBD3^* embryos. Quantification of the percentage of **(C)** total *Cen* mRNA and **(D)** mRNA in granules within the PCM zone (≤0.5 µm from γ-Tub surface). Each dot represents a measurement from 23 *Egl^WT^* and 39 *Egl^RBD3^* embryos. **(D’)** Log transformed RNA granule area from N=2304 granules from n=23 *Egl^WT^* embryos and N=4924 granules from n=39 *Egl^RBD3^* embryos; each dot represents a single granule. Mean ± SD displayed (red). Significance by unpaired t-test with **, P<0.01; and *** P<0.001. Scale bar: 5µm; 1µm (insets). (**E**) Model of the co-translational transport of *Cen* mRNA to centrosomes. Upon translation, an N’-terminal DLIC-binding CC1 box motif is exposed on the Cen nascent peptide, which associates with the dynein motor complex. We speculate Egl may bind to *Cen* mRNA. The Cen transport complex transits along microtubules via dynein to the centrosome. Additional on-site translation (Bergalet et al., 2020) may contribute to granule formation and/or stabilization.

## Discussion

While RNA localization to centrosomes is a longstanding observation, how local RNAs affect centrosome behavior remains relatively unstudied. That the localization of some centrosomal RNAs is conserved across taxa strongly implies a functional role. *Cen* mRNA serves as a valuable model to study this paradigm, as mislocalizing *Cen* mRNA leads to centrosome defects (Ryder et al., 2020). Further, the *Cen* 3’-UTR is important for targeting the antisense *ik2* mRNA, which codes for an actin regulatory factor (Oshima et al., 2006; Bergalet et al., 2020). Here, we examined what directs *Cen* mRNA to centrosomes.

We found the accumulation of *Cen* mRNA at centrosomes is puromycin-sensitive, highlighting the relevance of the nascent peptide for RNA localization. We then mapped domains within the Cen protein structure that enable RNA localization. Unexpectedly, we found *Cen* mRNA and protein localization can be separated. While the Cen N’-terminus is necessary but not sufficient for RNA localization, it is sufficient for Cen protein accumulation at centrosomes. These results argue for the presence of multiple domains that function cooperatively to target *Cen* mRNA to centrosomes.

We further defined the first 100 AA as important for *Cen* mRNA and protein localization. Within this region, we uncovered a conserved CC1 box that contributes to RNA localization.

Nevertheless, it is feasible that neighboring sequences also contribute to dynein or dynactin association, as shown for BicDR1 (Chaaban and Carter, 2022). CC1 boxes are found within dynein activating cargo adaptors, which directly bind DLIC and tether cargoes to the dynein motor complex (Gama et al., 2017; Lee et al., 2018; Lee et al., 2020; Chaaban and Carter, 2022). Dynein cargo adaptors also recruit the multi-subunit dynactin complex to the homodimeric Dhc, which enables dynein to translocate along the microtubule over long distances in association with cargo (Reck-Peterson et al., 2018). Known CC1 box-containing dynein activating cargo adaptors include the BicD, BicDR, and Spindly proteins. Our study positions Cen within this protein family. To support this, we show Cen biochemically associates with DLIC and that mutation of the CC1 box or various components of the dynein complex compromises *Cen* mRNA localization. The conservation of the CC1 box within CDR2 and CDR2L further suggests these mammalian proteins similarly function as dynein adaptors.

We therefore propose a model wherein the dynein complex directly binds the Cen CC1 box and, potentially, neighboring protein sequences as they emerge from the ribosome, activating the dynein motor complex, and directing Cen protein and mRNA to the centrosome (Figure 10E). While it is presently unknown if Egl directly binds *Cen* mRNA, it is tempting to speculate that such an interaction within the 3’-portion of *Cen* mRNA may be why this region of the *Cen* CDS is necessary for its localization. Thus, our model proposes that *Cen* mRNA localization requires both protein recognition via DLIC and RNA recognition by Egl. Such interactions are predicted to add valency and, therefore, robustness to the Cen transport complex. Of course, future work is needed to fully validate such a model.

Intriguingly, a recent study identified the RNAs for most *Drosophila* orthologs of the dynein cargo adaptors are enriched at the apical domain of follicle cell epithelia, where microtubules are nucleated. Further, the localization of those mRNAs was also translation-dependent (Cassella and Ephrussi, 2022). Similar observations were also noted for the centrosomal localization of *NIN* and *BICD2* mRNAs in mammalian cells (Safieddine et al., 2021) and parallel our findings of *Cen* mRNA in the syncytial embryo. Taken together, these findings strongly suggest that the translation-and dynein-dependent localization of dynein cargo adaptor RNAs to microtubule-organizing centers is a conserved feature.

For many RNAs, the assembly into higher order granules generally represents a translationally repressed state (Das et al., 2021). Paradoxically, the size of the *Cen* RNP seems to scale with Cen protein levels, as loss of the translational repressor FMRP (or *Cen* over-expression) enlarges the granule (Ryder et al., 2020). Moreover, we found that treatment with the translational inhibitor puromycin led to the rapid dissolution of the *Cen* RNP and depletion of its RNA at centrosomes. Thus, active translation not only directs *Cen* mRNA localization – it is required to maintain it. These findings are consistent with a requirement for continuous active transport or anchoring of *Cen* mRNA to the centrosome. Live imaging is required to distinguish these models and to directly visualize Cen translation en route to the centrosome, or within the granule itself.

## Materials and methods

### Fly stocks

The following *Drosophila* strains and transgenic lines were used: *y^1^w^1118^* (Bloomington *Drosophila* Stock Center, BDSC #1495) was the WT control; *P_BAC_-GFP-Cnn*, expressing Cnn tagged at the N-terminus with EGFP under endogenous regulatory elements (Lerit et al., 2015); *Ubi-GFP-γ-Tub23C,* expressing *GFP-γ-Tub* under the Ubiquitin promotor (Lerit and Rusan, 2013); Dynein heavy chain (*Dhc64C* gene) *Dhc^LOA^* is a hypomorphic allele that codes for F597Y mutant Dhc (modeled after the murine *Dync1h1* F580Y mutation (Salvador-Garcia et al., 2023); *Ubi*-*GFP-Dlic* (Pandey et al., 2007); *Cen^f04787^* is defined by a PiggyBac insertion in the *Cen* coding sequence and is a null mutation (BDSC #18805) (Bellen et al., 2004; Kao and Megraw, 2009). The *maternal α-Tub* promoter was used to drive *GAL4* expression (*matGAL4*; BDSC #7063) of *pUASp-Egl WT-FLAG-Egl shRNA* and *pUASp-Egl RBD3-FLAG-Egl shRNA* (Goldman et al., 2021), which were generous gifts from G. Gonsalvez (Augusta University), *Khc^RNAi^*(BDSC #35409; TRiP GL00330). Reduction of Khc by this TRiP line was previously demonstrated by western blot (Veeranan-Karmegam et al., 2016). *UASp-CenΔC*, *UASp-CenΔN, UASp-Cen^FL^-HA,* and *UASp-Cen^-ATG^-HA* were generated for this study (see below), expressed in the *Cen* null background, and driven with a single copy of *matGAL4*. To examine maternal effects, syncytial stage embryos were derived from mutant and/or transgenic mothers.

Flies were raised on cornmeal-based *Drosophila* medium (Bloomington formulation; Lab-Express, Inc.), and crosses were maintained at 25°C in a light and temperature-controlled chamber.

### Construction of transgenic and mutant animals

#### Cen CRISPR mutants

Strains with deletions within the region of the genome encoding the CC1 box of Cen were generated with the CRISPR/Cas9 reagents generated by the CRISPR Fly Design project (Port et al., 2014). A pCFD3 plasmid expressing a gRNA targeting the *Cen* sequence 5’-TACAATTGGCAGCAGAGCT-3’ (pCFD3-*Cen*) was generated by annealing overlapping oligos and ligating them with a Bbs1-cut backbone, as described previously (Port et al., 2014). The *Cen^Δ5^* allele was generated in an experiment in which a 100 ng/μl solution of pCFD3-*cen* was injected into *nos-cas9* embryos (CFD2 strain; (Port et al., 2014)). The *Cen^Δ12^*allele was generated in an experiment in which embryos were generated by crossing *nos-cas9* CFD2 females with males that had a stable integration of pCFD3-*Cen* at the attP40 docking site. In both these experiments, embryos were injected with a 150 ng/μl of a donor oligonucleotide (Ultramer; IDT) that contained codon changes that would, after precise homology-directed repair (HDR) of the Cas9 cleavage site, change A^25^ and A^26^ residues within the CC1 box to V residues. Previous work has shown that the equivalent mutation strongly reduces binding of cargo adaptors to dynein and dynactin (Schlager et al., 2014). The target site was amplified by PCR from genomic DNA extracted from the offspring of flies that developed from the embryos and analyzed by Sanger sequencing (Port and Bullock, 2016). Whilst the mutations carried by the donor oligonucleotide were not recovered, indicating that HDR was not successful, the *Δ12* allele was found.

#### Cen truncation lines

To generate the Cen truncations, the *Cen* coding sequence was divided into two pieces after the 289^th^ amino acid and PCR amplified using Phusion High-Fidelity DNA Polymerase from the cDNA clone pOT-LD41224 (*Drosophila* Genomics Resource Center (DGRC)). This site was chosen because it does not disrupt predicted secondary structure motifs. The following primers were used to amplify the respective pieces:

Cen N-terminal piece:

Forward: 5’-GGAAGTGGTGGTAGTGGAGGAAGTGAGGAATCCAATCACGGTTCGG-3’

Reverse 3’-TCGGCGCGCCCACCCTTTTAATCCCTCAGGCAGCGACT-5’

Cen C-terminal piece:

Forward: 5’-GAAGTGGTGGTAGTGGAGGAAGTATTAACGAAAGCAACACCAATATGGA-3’

Reverse: 5’-GGCGCGCCCACCCTTTTACTTTTGACGAAACTGATGATGATGAC-3’

Each Cen truncation was ligated into the pENTR-D Gateway vector (Invitrogen) using Gibson Assembly. The following primers and Phusion PCR were used to linearize the pENTR-D vector and add overlapping ends for ligation:

Vector for Cen N-terminal piece:

Forward: 5’-GTAGTCGCTGCCTGAGGGATTAAAAGGGTGGGCGCGC-3’

Reverse: 5’-CGCATAGTCAGGAACATCGTATGGGTACATGGTGAAGGGGGCGGC-3’

Vector for Cen C-terminal piece:

Forward: 5’-CAAGAGTCATCATCATCAGTTTCGTCAAAAGTAAAAGGGTGGGCGCGC-3’

Reverse: 5’-CGCATAGTCAGGAACATCGTATGGGTACATGGTGAAGGGGGCGGC-3’

A 3x HA tag plus linker was also incorporated by Gibson Assembly as a premade oligo with the following sequence. The 3x HA tag is underlined:

N-terminal 3x HA tag plus linker:

5’-ATGTACCCATACGATGTTCCTGACTATGCGGGCTATCCCTATGACGTCCCGGACTATGCAGGATCCTATCCATATGACGTTCCAGATTACGCTGGCGGCAGCGGTGGAAGTGGTGGTAGTGGAGGAAGT-3’

#### Cen FL and -ATG HA-tagged lines

To generate the full-length *Cen* HA-tagged construct, the full *Cen* CDS (including the ATG codon) was amplified using Phusion from pOT-LD41224 (DGRC) using the following primers: Full length *Cen* plus ATG:

Forward: 5’-GGCCGCCCCCTTCACCATGGAGGAATCCAATCACGGTTC-3’

Reverse: 5’-TCCACCGCTGCCGCCCTTTTGACGAAACTGATGATGATGAC-3’

The pENTR-D vector was linearized by Phusion PCR using the following primers:

Forward: 5’-CCTATCCATATGACGTTCCAGATTACGCTTAAAAGGGTGGGCGCGCC-3’

Reverse: 5’-CGAACCGTGATTGGATTCCTCCATGGTGAAGGGGGCGGC-3’

The full-length *Cen* was inserted into the pENTR-D vector with a C-terminal 3x HA tag plus linker by Gibson assembly. The C-terminal 3x HA tag plus linker was added as a premade oligo with the following sequence. The 3x HA tag is underlined:

C-terminal 3x HA tag plus linker:

5’-GGCGGCAGCGGTGGAAGTGGTGGTAGTGGAGGAAGTTACCCATACGATGTTCCTGACTATGCGGGCTATCCCTATGACGTCCCGGACTATGCAGGATCCTATCCATATGACGTTCCAGATTACGCTTAA-3’

To generate the *Cen* -ATG HA-tagged construct, the *Cen* coding sequence was amplified from pOT-LD41224 using the following primers to remove the initiating ATG codon then ligated by Gibson Assembly, as described for the full-length construct:

Forward: 5’-CCGCGGCCGCCCCCTTCACCGAGGAATCCAATCACGGTTC-3’

Reverse: 5’-CCGTGATTGGATTCCTCGGTGAAGGGGGCGGC-3’

For all lines, sequence-verified single colony clones were shuttled into the destination vector *pPWattB* (UASp-Gateway with attB sites for Phi31C transformation) using the Gateway cloning system (Invitrogen). Constructs were inserted at the *attP2* (Chromosome III) locus and transgenic animals were generated by BestGene, Inc.

### Sequence alignment

Protein sequences were obtained from UniProt (UniProt, 2023), aligned using the Clustal Omega multiple sequence alignment tool (Madeira et al., 2022), then displayed using ESPript 3.0 (Robert and Gouet, 2014) using the percent equivalent similarity and the black-and-white color schemes.

### Immunofluorescence

Embryos (0.5–2.5 hr) were collected then dechorionated, fixed in 4% paraformaldehyde, and blocked in BBT (PBS supplemented with 0.1% Tween-20 and 0.1% BSA) as described in (Lerit et al., 2015). Primary antibodies were diluted in BBT in incubated done overnight at 4 °C with nutation. On the following day, samples were washed three times with BBT then blocked again with BBT supplemented with 2% normal goat serum (NGS) prior to incubation with secondary antibodies and DAPI for 2 hours at room temperature. Samples were mounted in AquaPoly/Mount (VWR, 87001-902).

To visualize microtubules, embryos were fixed in a solution of 1:1 of heptane:37% formaldehyde for 3 min with intermittent mixing, then manually devitellinized using 30G PrecisionGlide needles (BD) (Theurkauf, 1994). Embryos were then blocked in BBT, rinsed in PBS, blocked again with Image-iT FX signal enhancer (ThermoScientific), then incubated with antibodies overnight at 4 °C with primary antibodies diluted in BBT. On the following day, samples were processed and mounted as described above.

The following antibodies were used: rabbit anti-Cen (UT393, 1:500; gift from T. Megraw, Florida State University) (Kao and Megraw, 2009) recognizes the C-terminus of Cen, rabbit anti-Cen (1:500; gift from T. Megraw) (Kao and Megraw, 2009) recognizes the N-terminus of Cen, mouse anti-α-Tub DM1a (1:500; Sigma, T6199), and rabbit anti-Cnn (1:4000; gift from T. Megraw). Secondary antibodies and stains: Alexa Fluor 488, 568, or 647 (1:500, Invitrogen). DAPI was used at 10 ng/mL (Thermo Fisher).

### Single molecule fluorescence *in situ* hybridization (smFISH)

smFISH experiments were conducted as previously described in (Ryder et al., 2020). All steps were done using RNase-free solutions. In short, 0–2-hour embryos were aged 30 minutes then fixed in 4% paraformaldehyde and rehydrated stepwise into 0.1% PBST. Rehydrated embryos were then washed with wash buffer (WB; 10% formamide and 2× SSC supplemented fresh each experiment with 0.1% Tween-20 and 2 μg/mL nuclease-free BSA) at room temperature and incubated in a freshly made hybridization buffer (HB; 100 mg/mL dextran sulfate and 10% formamide in 2× SSC supplemented fresh each experiment with 0.1% Tween-20, 2 μg/mL nuclease-free BSA and 10 mM ribonucleoside vanadyl complex (RVC; S1402S; New England Biolabs)) in a 37 °C water bath. Embryos were incubated overnight in a 37 °C water bath in HB with a final concentration of 0.4 µM Stellaris smFISH probes (*Cen* or *GAPDH*) conjugated to Quasar 570 dye (LGC Biosearch Technologies). The following day, the hybridized embryos were washed with WB three times, then with 0.1% PBST, and stained with DAPI (1:1000). Vectashield mounting medium (Vector Laboratories, H-1000) was used to mount the slides. Complete probe sequences are reported in Ryder et al., 2020.

### Dual smFISH and immunofluorescence

Dual smFISH and IF experiments were as previously described (Ryder et al., 2020; Fang and Lerit, 2022). All steps were done using RNase-free solutions. Embryos were rehydrated and washed first in 0.1% PBST (PBS plus 0.1% Tween-20) and then in WB, as above. Embryos were then incubated with 100 µL of HB for 10-20 minutes in a 37 °C water bath, followed by an overnight incubation in 25 µL of HB containing 0.5 μM smFISH probes and primary antibody in a 37 °C water bath. On the next day, embryos were washed four times for 30 minutes in prewarmed WB, stained with secondary antibody and DAPI (1:1000) for 2 hours at room temperature, washed with 0.1% PBST, and mounted with Vectashield mounting medium (H-1000; Vector Laboratories). Slides were stored at 4 °C and imaged within 1 week.

### Pharmacological inhibition of translation

0.5–2.5 hr embryos were collected and incubated in a 1:1 solution (450 µL: 450 µL) of heptane: Robb’s medium (1 mM calcium chloride, 10 mM glucose, 100 mM HEPES (pH 7.2), 1.2 mM MgCl_2_, 55 mM KOAc, 40 mM NaOAc, and 100 mM sucrose) containing the appropriate drug or an equivalent volume of DMSO. The concentrations and incubation times for each drug were: 3 mM puromycin (Sigma-Aldrich P8833) for 10 min; 0.1 mM anisomycin (Sigma-Aldrich A9789) for 15 min; and 0.71 mM cycloheximide (VWR, 97064-724) for 15 min. After drug incubation, Robb’s medium was removed, and 450 µl of 4% paraformaldehyde in PBS was added, and embryos were fixed for 20 min, then devitellinized. Samples were processed for smFISH or dual smFISH + IF, as above.

### Microtubule regrowth assay

For cold-shock, embryos were transferred to a 1.5 mL tube and incubated on ice for 5 minutes to disrupt the microtubules, then immediately fixed. For microtubule regrowth (recovery), cold-shocked embryos were incubated in room-temperature PBS for 5 minutes, then immediately fixed. Control, cold-shocked, and recovery embryos were then processed for sequential smFISH and immunofluorescence, as follows. Embryos were fixed in 37% formaldehyde and manually devitellinized, as described above, rinsed in 0.1% PBST, incubated in Image-IT FX for 30 minutes, washed again in 0.1% PBST, and then washed in WB buffer for 10 minutes. Embryos were incubated in HB buffer for 10-20 minutes in a 37 °C water bath prior to an overnight incubation in 25 μL of HB containing 0.5 μM smFISH probes. On the next day, embryos were washed in WB buffer, 2 X SSC with 0.1% Tween-20, and then 0.1% PBST sequentially. Next, embryos were blocked in 0.1% BBT buffer (PBS supplemented with 0.1% BSA and 0.1% Tween-20). Embryos were then incubated overnight at 4 °C with primary antibody in 0.1% BBT, further blocked in 0.1% BBT supplemented with 2% NGS, and incubated for 2 hours at room temperature with secondary antibodies and DAPI. Embryos were mounted in Vectashield mounting medium prior to imaging. Slides were stored at 4 °C and imaged within 1 week.

### Microscopy

Images were acquired on a Nikon Ti-E system fitted with a Yokogawa CSU-X1 spinning disk head (Yokogawa Corporation of America), Orca Flash 4.0 v2 digital complementary metal-oxide-semiconductor camera (Hamamatsu Corporation), Perfect Focus system (Nikon), and a Nikon LU-N4 solid state laser launch (15 mW 405, 488, 561, and 647 nm) using the following objectives: 100x 1.49 NA Apo TIRF oil immersion or 40x 1.3 NA Plan Fluor oil immersion. Images were acquired at ambient temperature (∼25°C) using either Vectashield or Aqua-Poly/Mount imaging medium, as described, using Nikon Elements AR software.

### Image Analysis

Images for figures were assembled using Fiji (NIH; (Schindelin et al., 2012)) and Adobe Illustrator. The software was used to separate or merge channels, crop regions of interest, generate maximum intensity projections, and adjust brightness and contrast.

#### RNA detection and measurements

Raw, single channel .tif files of centrosomes and RNA were segmented in three dimensions using a code adapted from the Allen Institute for Cell Science Cell Segmenter then run through the open-source, Python-based SubcellularDistribution pipeline (Ryder and Lerit, 2020) to calculate the percentage of RNA overlapping with centrosomes, percent of RNA in granules, and granular intensities. Granules at a distance 0µm or 0.5µm with a normalized intensity greater than 4 were log transformed and plotted using R. Unless otherwise noted, all RNA measurements were calculated based on the percentage of *Cen* mRNA residing within 0 µm (i.e., overlapping) from the Cnn surface.

We examined *Cen* mRNA distributions in *Egl^WT^* versus *Egl^RBD3^* embryos by smFISH relative to the PCM marker γ-Tub-GFP rather than GFP-Cnn because the *GFP-Cnn* and *Egl* transgenes both reside on Chromosome III. Because γ-Tub occupies a significantly smaller radius of the PCM than Cnn (about 600 vs 1400 nm, by structured illumination microscopy; (Lerit et al., 2015)), for these experiments, we took a conservative measurement of the percentage of *Cen* mRNA residing within 0.5 µm from the γ-Tub surface.

Because cold-shock compresses the volume of Cnn, for the microtubule regrowth experiments, the percentage of *Cen* mRNA residing within 1 µm from the surface of the core centriolar protein, Asl, was measured from all samples (control, cold-shock, and recovery).

#### Spindle morphology defects

To quantify spindle morphology, mitotic embryos imaged at 40x were examined for the following morphologies: bent spindles, multipolar or fused spindles, acentrosomal spindle poles, and defective centrosome separation. If any spindles within an embryo contained one of these phenotypes, the embryo was considered positive for a spindle morphology defect. Three independent biological replicates were performed for each genotype.

#### Colocalization analysis

Single optical slices were analyzed for co-occurrence of *Cen* RNA with microtubules. Mander’s M1 coefficient was calculated from a 66.56 µm^2^ area using the JacoP plugin for ImageJ in which a manual threshold was applied to remove background signal (Bolte and Cordelieres, 2006).

### Immunoprecipitation

To examine the interaction between DLIC and Cen, the immunoprecipitation was performed as in (Dix et al., 2013) using 0–5 hour embryos lysed in a buffer containing 25 mM HEPES pH 7.0, 50 mm KCl, 1 mM MgCl2, 2 mM DTT and 250 mM sucrose, supplemented with 1x Complete protease inhibitors (Roche). Transgenic strains expressing GFP-Dlic (Pandey et al., 2007) were used for immunoprecipitation using agarose GFP-Trap beads (Chromotek).

To determine the amino acid residues corresponding to the truncated Cen^-ATG^ protein product, we immunoprecipitated Cen from ovaries harvested from well-fed 1–2-day old *Cen* null females expressing *Cen^FL^* or *Cen^-ATG^* transgenes lysed in RCB buffer containing 50 mM HEPES, pH 7.4, 150 mM NaCl, 2.5 mM MgCl_2_, 0.01% Triton x-100, and 250 mM sucrose supplemented with 1x Complete protease inhibitors, 1 mM DTT, and 1 µg/mL Pepstatin A. Protein concentration was normalized across samples using a Pierce BCA assay (Thermo Scientific, cat. 23225). We used ∼50-pairs of ovaries, yielding about 10 mg of protein per reaction. For each reaction, 50 µL of Pierce Protein A/G magnetic beads (Thermo Scientific, cat. 88802) were prewashed in RCB, and half of the bead slurry was used to preclear the lysate for 30 min to minimize nonspecific binding. The precleared lysate was reserved and incubated with 10 µL of a 1:50 dilution of rabbit anti-HA antibody (C29F4, Cell Signaling Technology) for 2-hr at RT. The remaining 25 µL of prewashed beads was then added and incubated for 1-hr at RT. Beads were washed well in RCB, then processed for immunoblotting (10 µg protein per lane) or shipped to MS Bioworks for mass spectrometry (see below).

### Mass Spectrometry

Mass spectrometry and analysis was performed by MS Bioworks, LLC (Ann Arbor, MI). 3 x 20 μL per immunoprecipitated Cen-HA sample were separated on a 4-12% Bis-Tris NuPAGE Novex mini-gel (Invitrogen) using the MOPS buffer system. The gel was stained with Coomassie, and target bands excised. Gel segments were digested with three enzymes using a robot (DigestPro, CEM) with the following protocol. First, they were washed with 25 mM ammonium bicarbonate followed by acetonitrile. Next, samples were reduced with 10 mM DTT at 60°C followed by alkylation with 50mM iodoacetamide at RT. Next, samples were digested with trypsin/ chymotrypsin/ elastase (Promega) at 37°C for 4h. The reaction was quenched with formic acid and the supernatant was analyzed directly without further processing. The gel digests were analyzed by nano LC/MS/MS with a Waters M-class HPLC system interfaced to a ThermoFisher Fusion Lumos. Peptides were loaded on a trapping column and eluted over a 75 μm analytical column at 350nL/min; both columns were packed with XSelect CSH C18 resin (Waters). A 30 min gradient was employed. The mass spectrometer was operated in data-dependent mode, with MS and MS/MS performed in the Orbitrap at 60,000 FWHM resolution and 15,000 FWHM resolution, respectively. APD was turned on. The instrument was run with a 3s cycle for MS and MS/MS. From the FL immunoprecipitation, 152 proteins were identified, and 4703 spectra were matched. Cen was the second-most abundant protein. From the -ATG immunoprecipitation, 302 proteins were identified, and 9706 spectra were matched. Cen was the fourth-most abundant protein.

### Immunoblotting

Western blotting was performed as in (Dix et al., 2013). Alternatively, appropriately aged embryos were lysed in 0.1% PBST using an electric homogenizer on ice then immediately boiled in 5x SDS loading dye for 5 minutes and returned to ice. Samples were resolved by premade gradient SDS-PAGE gel (Bio-Rad) and transferred to nitrocellulose membrane by wet or semi-dry transfer. Membranes were blocked in a 5% dry milk solution diluted in TBST (Tris-based saline with 0.05% Tween-20) and incubated overnight at 4°C with primary antibodies in 1% dry milk in TBST solution. Primary antibodies used: rabbit anti-C terminus of Cen (UT393) or rabbit anti-N terminus of Cen (both 1:5000; gift from T. Megraw, Florida State University, (Kao and Megraw, 2009)), guinea pig anti-Asterless (1:5000; gift from G. Rogers, University of Arizona), mouse anti-actin JLA20-S (1:1000; DSHB), mouse anti-β-Tub E7 (1:15,000; DSHB), rabbit anti-HA (1: 5000; C29F4, Cell Signaling Technology), mouse anti-BicD 1B11 (1:1000; (Suter and Steward, 1991)), and mouse anti-GFP clones 7.1 and 13.1 (1:1000; Roche).

The following day, membranes were washed with TBST and incubated with secondary antibodies for 1 hour at room temperature. Secondary antibodies were diluted 1:2500 in TBST and included goat anti-mouse HRP (31430; Thermo Fisher Scientific), goat anti-rabbit HRP (31460, ThermoFisher Scientific), and goat anti-guinea pig HRP (A18769, ThermoFisher Scientific). Bands were visualized with Clarity ECL substrate (1705061; Bio-Rad) on a Bio-Rad ChemiDoc imaging system.

### qPCR

Embryos aged 0.5–2.5 hours were dechorionated with bleach, flash frozen in liquid nitrogen, and stored at -80 °C. A volume of 100 µL of embryos per biological replicate was homogenized in TRIzol (Invitrogen) and RNA was extracted using phenol: chloroform extraction. The extracted RNA was then treated with Ambion TURBO DNase (Thermo Fisher Scientific, AM2238). cDNA was then synthesized using an iScript cDNA synthesis kit (Bio-Rad, 170-8891).

Three technical replicates per biological replicate were run concurrently in a 96-well plate (Bio-Rad, HSP9601) using iQ SYBR Green Supermix (Bio-Rad, 170-8882). Data was collected on a Bio-Rad CFX96 Real-time machine. Levels of *Cen* expression were normalized to Ribosomal protein L32 (RP49). The following primers were used:

*RP49* (amplicon 75 base pairs)

Forward: 5’-CATACAGGCCCAAGATCGTG-3’

Reverse: 5’-ACAGCTTAGCATATCGATCCG-3’

*Centrocortin* (amplicon 78 base pairs)

Forward: 5’-AAAGTACCCCCGGTAACACC-3’

Reverse: 5’-TGAGGATACGACGCTCTGTG-3’

### Statistical Analysis

All statistical analyses were conducted using GraphPad Prism software, except granule area was calculated using R-software. Data were first tested for any outliers using a ROUT test with a Q= 1% and normality using the D’Agostino and Pearson normality test. This was followed by Student’s two-tailed t test, ANOVA, Fisher’s exact test, or the appropriate nonparametric tests. Data were plotted with mean ± SD displayed.

## Supplemental material

Two supplemental figures accompany this study.

**Supplemental Figure 1.**
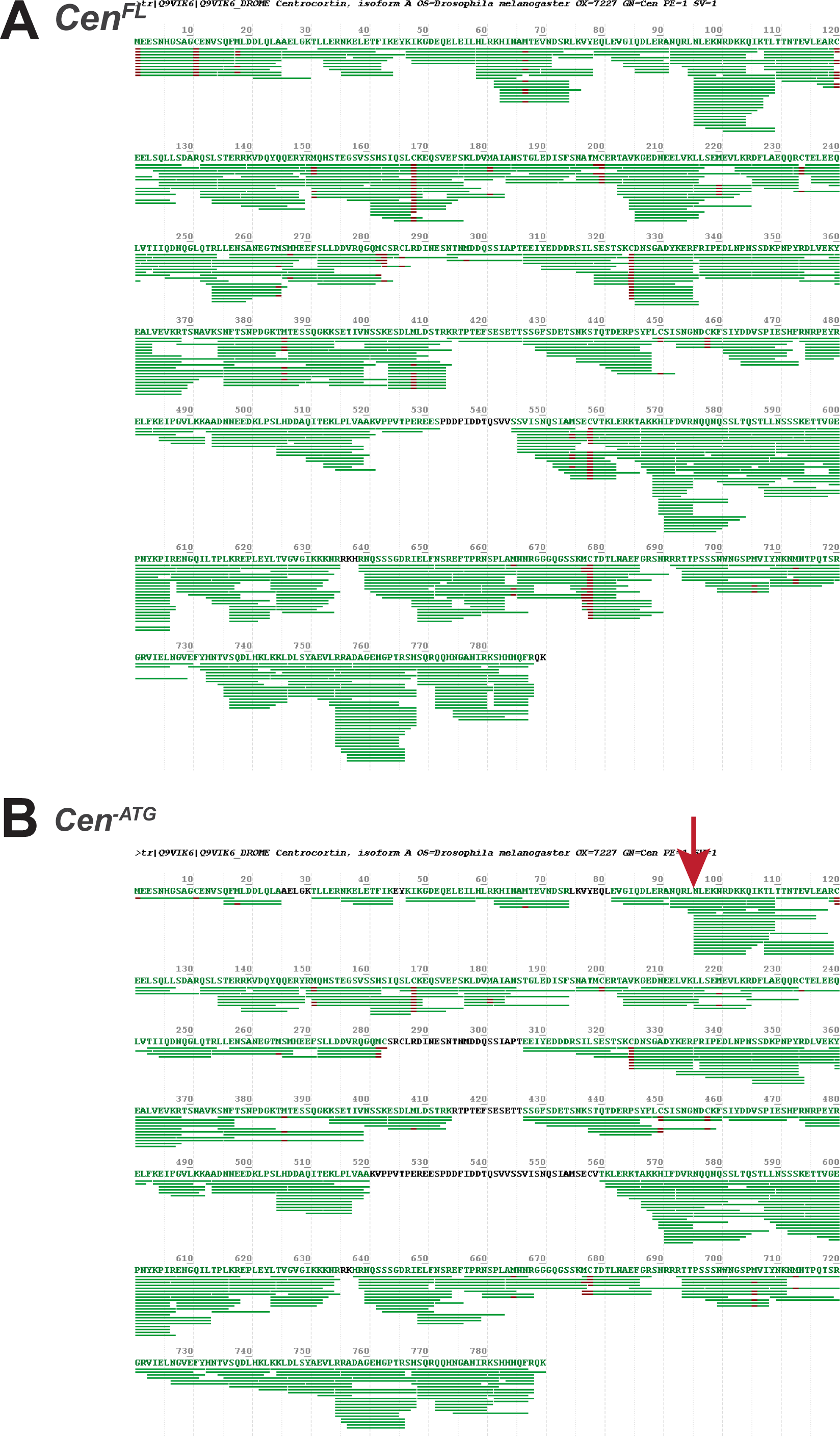
Mass spectrometry analysis of Cen protein products. **(B)** Sequence mapping of spectra (green lines) from (A*) Cen^FL^* and (B) *Cen^-ATG^*, as identified by mass spectrometry following anti-HA immunoprecipitation from 1–2-day ovarian extracts. The HA-tagged constructs were expressed in the *Cen* null background. The UniProt reference Cen sequence used was Q9VIK6. The arrow marks the most N’-terminal position where abundant Cen spectra map to the *Cen^-ATG^* protein product. The first 90–100 AA are not well covered by the spectra and are likely absent from the truncation. Data shown are representative of two independent experiments.

**Supplemental Figure 2.**
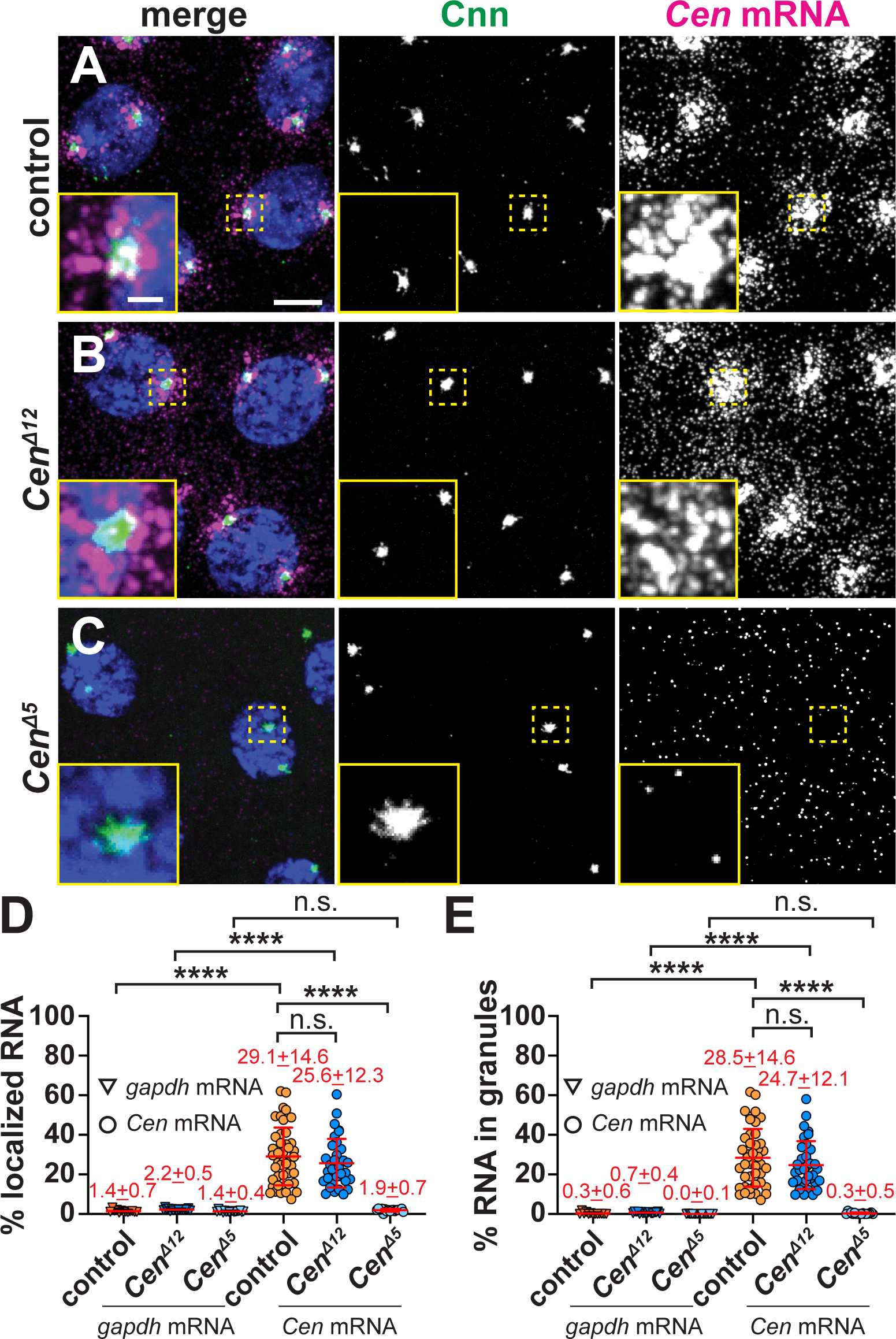
*Cen* mRNA localization in early embryos. Maximum-intensity projections of NC 11 interphase embryos expressing *GFP-Cnn* (green) stained with C*en* smFISH (magenta) and DAPI (blue nuclei). **(A)** Control embryos show C*en* mRNA enriched at centrosomes, primarily in RNPs, which are also present in **(B)** *Cen^Δ12^* samples. **(C)** *Cen* mRNA localization and granule formation are severely impaired in *Cen^Δ5^* embryos. Quantification of the percentage of *Cen* or *gapdh* mRNAs **(D)** overlapping with the centrosome surface and **(E)** residing in granules (0 µm distance from Cnn). Each dot represents a single measurement from control (N= 13 *gapdh* and 44 *Cen* mRNA), *Cen^Δ12^* (N= 17 *gapdh* and 34 *Cen* mRNA), and *Cen^Δ5^*(N= 18 *gapdh* and 17 *Cen* mRNA) labelled embryos. Mean ± SD shown. ****, P<0.0001 by Brown-Forsythe and Welch ANOVA tests followed by Dunnett’s T3 multiple comparison test. Scale bar: 5µm; 1µm (insets).

## Data Availability Statement

All data are available in the published article and its online supplemental material. Source files of uncropped versions of the immunoblots presented in the figures are included.

## Acknowledgments

We thank Drs. Nasser Rusan, Greg Rogers, Timothy Megraw, Jordan Raff, and Graydon Gonsalvez for gifts of reagents and Drs. Gary Bassell and Sulagna Das for critical reading of this manuscript. We acknowledge Dr. Richard Jones from MS Bioworks, LLC for technical expertise and assistance with mass spectrometry and analysis. We are also grateful to Dr. Pearl Ryder for early contributions to this work. Stocks obtained from the Bloomington *Drosophila* Stock Center (NIH grant P40OD018537); antibodies from the Developmental Studies Hybridoma Bank, created by the NICHD of the NIH and maintained at the University of Iowa Department of Biology; and reagents from the *Drosophila* Genomics Resource Center (DGRC), supported by NIH grant 2P40OD010949, were used in this study.

This work was supported by NIH grants 5K12GM000680 (HZ-S), 1F31NS134380 (JB), 5K99GM143517 (JF), and R01GM138544 (DAL); the UK Medical Research Council (as part of United Kingdom Research and Innovation (file reference number MC_U105178790) (SLB); and the NIH administrative supplements 3R01GM138544-01S1 (for JL) and 3R01GM138544-03S1 (for CAH). DAL is also supported by a Research Scholar Grant (RSG-22-874157-01-CCB) from the American Cancer Society.

The authors declare no competing financial interests.

## Author Contributions

**H. Zein-Sabatto**– Conceived of and designed the experiments, generated reagents, performed the experiments, analyzed the data, made the figures, secured funding, supervised the project, wrote the original draft of the manuscript, and edited the manuscript. **J. Brockett**– Performed the experiments and analyzed the data. **L. Jin**– Generated reagents and performed the experiments. **C.A. Husbands**– Generated reagents, performed the experiments, and analyzed the data. **J. Lee**– Generated reagents, performed the experiments, and analyzed the data. **J. Fang**– Performed the experiments and analyzed the data. **J. Buehler**– Analyzed the data. **S.L. Bullock**– Conceived of and designed the experiments, performed the experiments, secured funding, supervised the project, wrote the original draft of the manuscript, and edited the manuscript. **D.A. Lerit**– Conceived of and designed the experiments, made the figures, secured funding, supervised the project, wrote the original draft of the manuscript, and edited the manuscript.

All authors approved of the manuscript.

## Acknowledgments

αTub: α-Tubulin
Aniso: Anisomycin
Asl: Asterless
BicD: Bicaudal
CDR2: Cerebellar degeneration-related protein 2
CDR2L: Cerebellar degeneration-related protein 2-like
Cen: Centrocortin
CHX: Cycloheximide
Cnn: Centrosomin
Dhc: dynein heavy chain
DLIC: dynein light intermediate chain
Egl: Egalitarian
γTub: γ-Tubulin
Khc: kinesin heavy chain
NC: nuclear cycle
PCM: pericentriolar material
Puro: puromycin
RBD: RNA-binding domain

## Notes

### Competing Interest Statement

The authors have declared no competing interest.

## References

Apweiler, R., T.K. Attwood, A. Bairoch, A. Bateman, E. Birney, M. Biswas, P. Bucher, L. Cerutti, F. Corpet, M.D. Croning, R. Durbin, L. Falquet, W. Fleischmann, J. Gouzy, H. Hermjakob, N. Hulo, I. Jonassen, D. Kahn, A. Kanapin, Y. Karavidopoulou, R. Lopez, B. Marx, N.J. Mulder, T.M. Oinn, M. Pagni, F. Servant, C.J. Sigrist, E.M. Zdobnov, and C. InterPro. 2000. InterPro--an integrated documentation resource for protein families, domains and functional sites. Bioinformatics. 16:1145–1150.

Bellen, H.J., R.W. Levis, G. Liao, Y. He, J.W. Carlson, G. Tsang, M. Evans-Holm, P.R. Hiesinger, K.L. Schulze, G.M. Rubin, R.A. Hoskins, and A.C. Spradling. 2004. The BDGP gene disruption project: single transposon insertions associated with 40% of Drosophila genes. Genetics. 167:761–781.

Bergalet, J., D. Patel, F. Legendre, C. Lapointe, L.P. Benoit Bouvrette, A. Chin, M. Blanchette, E. Kwon, and E. Lecuyer. 2020. Inter-dependent Centrosomal Co-localization of the cen and ik2 cis-Natural Antisense mRNAs in Drosophila. Cell Rep. 30:3339–3352 e3336.

Besse, F., and A. Ephrussi. 2008. Translational control of localized mRNAs: restricting protein synthesis in space and time. Nat Rev Mol Cell Biol. 9:971–980.

Bolte, S., and F.P. Cordelieres. 2006. A guided tour into subcellular colocalization analysis in light microscopy. J Microsc. 224:213–232.

Bullock, S.L. 2007. Translocation of mRNAs by molecular motors: think complex? Semin Cell Dev Biol. 18:194–201.

Cassella, L., and A. Ephrussi. 2022. Subcellular spatial transcriptomics identifies three mechanistically different classes of localizing RNAs. Nat Commun. 13:6355.

Chaaban, S., and A.P. Carter. 2022. Structure of dynein-dynactin on microtubules shows tandem adaptor binding. Nature. 610:212–216.

Chin, A., and E. Lecuyer. 2017. RNA localization: Making its way to the center stage. Biochim Biophys Acta Gen Subj. 1861:2956–2970.

Chouaib, R., A. Safieddine, X. Pichon, A. Imbert, O.S. Kwon, A. Samacoits, A.M. Traboulsi, M.C. Robert, N. Tsanov, E. Coleno, I. Poser, C. Zimmer, A. Hyman, H. Le Hir, K. Zibara, M. Peter, F. Mueller, T. Walter, and E. Bertrand. 2020. A Dual Protein-mRNA Localization Screen Reveals Compartmentalized Translation and Widespread Co-translational RNA Targeting. Dev Cell. 54:773–791 e775.

Das, S., M. Vera, V. Gandin, R.H. Singer, and E. Tutucci. 2021. Intracellular mRNA transport and localized translation. Nat Rev Mol Cell Biol. 22:483–504.

Dienstbier, M., F. Boehl, X. Li, and S.L. Bullock. 2009. Egalitarian is a selective RNA-binding protein linking mRNA localization signals to the dynein motor. Genes Dev. 23:1546–1558.

Dix, C.I., H.C. Soundararajan, N.S. Dzhindzhev, F. Begum, B. Suter, H. Ohkura, E. Stephens, and S.L. Bullock. 2013. Lissencephaly-1 promotes the recruitment of dynein and dynactin to transported mRNAs. J Cell Biol. 202:479–494.

Fang, J., and D.A. Lerit. 2022. Orb-dependent polyadenylation contributes to PLP expression and centrosome scaffold assembly. Development. 149.

Foe, V.E., and B.M. Alberts. 1983. Studies of nuclear and cytoplasmic behaviour during the five mitotic cycles that precede gastrulation in Drosophila embryogenesis. J Cell Sci. 61:31–70.

Gama, J.B., C. Pereira, P.A. Simoes, R. Celestino, R.M. Reis, D.J. Barbosa, H.R. Pires, C. Carvalho, J. Amorim, A.X. Carvalho, D.K. Cheerambathur, and R. Gassmann. 2017. Molecular mechanism of dynein recruitment to kinetochores by the Rod-Zw10-Zwilch complex and Spindly. J Cell Biol. 216:943–960.

Gasparski, A.N., D.E. Mason, K. Moissoglu, and S. Mili. 2022. Regulation and outcomes of localized RNA translation. Wiley Interdiscip Rev RNA. 13:e1721.

Gepner, J., M. Li, S. Ludmann, C. Kortas, K. Boylan, S.J. Iyadurai, M. McGrail, and T.S. Hays. 1996. Cytoplasmic dynein function is essential in Drosophila melanogaster. Genetics. 142:865–878.

Goldman, C.H., H. Neiswender, F. Baker, R. Veeranan-Karmegam, S. Misra, and G.B. Gonsalvez. 2021. Optimal RNA binding by Egalitarian, a Dynein cargo adaptor, is critical for maintaining oocyte fate in Drosophila. RNA Biol. 18:2376–2389.

Goldman, C.H., H. Neiswender, R. Veeranan-Karmegam, and G.B. Gonsalvez. 2019. The Egalitarian binding partners Dynein light chain and Bicaudal-D act sequentially to link mRNA to the Dynein motor. Development. 146.

Gould, R.R., and G.G. Borisy. 1977. The pericentriolar material in Chinese hamster ovary cells nucleates microtubule formation. J Cell Biol. 73:601–615.

Groisman, I., Y.S. Huang, R. Mendez, Q. Cao, W. Theurkauf, and J.D. Richter. 2000. CPEB, maskin, and cyclin B1 mRNA at the mitotic apparatus: implications for local translational control of cell division. Cell. 103:435–447.

Grollman, A.P. 1967. Inhibitors of protein biosynthesis. II. Mode of action of anisomycin. J Biol Chem. 242:3226–3233.

Hafezparast, M., R. Klocke, C. Ruhrberg, A. Marquardt, A. Ahmad-Annuar, S. Bowen, G. Lalli, A.S. Witherden, H. Hummerich, S. Nicholson, P.J. Morgan, R. Oozageer, J.V. Priestley, S. Averill, V.R. King, S. Ball, J. Peters, T. Toda, A. Yamamoto, Y. Hiraoka, M. Augustin, D. Korthaus, S. Wattler, P. Wabnitz, C. Dickneite, S. Lampel, F. Boehme, G. Peraus, A. Popp, M. Rudelius, J. Schlegel, H. Fuchs, M. Hrabe de Angelis, G. Schiavo, D.T. Shima, A.P. Russ, G. Stumm, J.E. Martin, and E.M. Fisher. 2003. Mutations in dynein link motor neuron degeneration to defects in retrograde transport. Science. 300:808–812.

Holt, C.E., and S.L. Bullock. 2009. Subcellular mRNA localization in animal cells and why it matters. Science. 326:1212–1216.

Jung, H., C.G. Gkogkas, N. Sonenberg, and C.E. Holt. 2014. Remote control of gene function by local translation. Cell. 157:26–40.

Kao, L.R., and T.L. Megraw. 2009. Centrocortin cooperates with centrosomin to organize Drosophila embryonic cleavage furrows. Curr Biol. 19:937–942.

Khodjakov, A., and C.L. Rieder. 1999. The sudden recruitment of gamma-tubulin to the centrosome at the onset of mitosis and its dynamic exchange throughout the cell cycle, do not require microtubules. J Cell Biol. 146:585–596.

Kislauskis, E.H., Z. Li, R.H. Singer, and K.L. Taneja. 1993. Isoform-specific 3’-untranslated sequences sort alpha-cardiac and beta-cytoplasmic actin messenger RNAs to different cytoplasmic compartments. J Cell Biol. 123:165–172.

Lecuyer, E., H. Yoshida, N. Parthasarathy, C. Alm, T. Babak, T. Cerovina, T.R. Hughes, P. Tomancak, and H.M. Krause. 2007. Global analysis of mRNA localization reveals a prominent role in organizing cellular architecture and function. Cell. 131:174–187.

Lee, I.G., S.E. Cason, S.S. Alqassim, E.L.F. Holzbaur, and R. Dominguez. 2020. A tunable LIC1-adaptor interaction modulates dynein activity in a cargo-specific manner. Nat Commun. 11:5695.

Lee, I.G., M.A. Olenick, M. Boczkowska, C. Franzini-Armstrong, E.L.F. Holzbaur, and R. Dominguez. 2018. A conserved interaction of the dynein light intermediate chain with dynein-dynactin effectors necessary for processivity. Nat Commun. 9:986.

Lerit, D.A. 2022. Signed, sealed, and delivered: RNA localization and translation at centrosomes. Mol Biol Cell. 33.

Lerit, D.A., H.A. Jordan, J.S. Poulton, C.J. Fagerstrom, B.J. Galletta, M. Peifer, and N.M. Rusan. 2015. Interphase centrosome organization by the PLP-Cnn scaffold is required for centrosome function. J Cell Biol. 210:79–97.

Lerit, D.A., and N.M. Rusan. 2013. PLP inhibits the activity of interphase centrosomes to ensure their proper segregation in stem cells. J Cell Biol. 202:1013–1022.

Macdonald, P.M., and G. Struhl. 1988. cis-acting sequences responsible for anterior localization of bicoid mRNA in Drosophila embryos. Nature. 336:595–598.

Mach, J.M., and R. Lehmann. 1997. An Egalitarian-BicaudalD complex is essential for oocyte specification and axis determination in Drosophila. Genes Dev. 11:423–435.

Madeira, F., M. Pearce, A.R.N. Tivey, P. Basutkar, J. Lee, O. Edbali, N. Madhusoodanan, A. Kolesnikov, and R. Lopez. 2022. Search and sequence analysis tools services from EMBL-EBI in 2022. Nucleic Acids Res. 50:W276–W279.

Marshall, W.F., and J.L. Rosenbaum. 2000. Are there nucleic acids in the centrosome? Curr Top Dev Biol. 49:187–205.

Martin, K.C., and A. Ephrussi. 2009. mRNA localization: gene expression in the spatial dimension. Cell. 136:719–730.

McClintock, M.A., C.I. Dix, C.M. Johnson, S.H. McLaughlin, R.J. Maizels, H.T. Hoang, and S.L. Bullock. 2018. RNA-directed activation of cytoplasmic dynein-1 in reconstituted transport RNPs. Elife. 7.

Megraw, T.L., S. Kilaru, F.R. Turner, and T.C. Kaufman. 2002. The centrosome is a dynamic structure that ejects PCM flares. J Cell Sci. 115:4707–4718.

Mitchison, T., and M. Kirschner. 1984. Microtubule assembly nucleated by isolated centrosomes. Nature. 312:232–237.

Mofatteh, M., and S.L. Bullock. 2017. SnapShot: Subcellular mRNA Localization. Cell. 169:178–178 e171.

Nathans, D. 1964. Puromycin Inhibition of Protein Synthesis: Incorporation of Puromycin into Peptide Chains. Proc Natl Acad Sci U S A. 51:585–592.

Navarro, C., H. Puthalakath, J.M. Adams, A. Strasser, and R. Lehmann. 2004. Egalitarian binds dynein light chain to establish oocyte polarity and maintain oocyte fate. Nat Cell Biol. 6:427-435.

Nolan, P.M., J. Peters, M. Strivens, D. Rogers, J. Hagan, N. Spurr, I.C. Gray, L. Vizor, D. Brooker, E. Whitehill, R. Washbourne, T. Hough, S. Greenaway, M. Hewitt, X. Liu, S. McCormack, K. Pickford, R. Selley, C. Wells, Z. Tymowska-Lalanne, P. Roby, P. Glenister, C. Thornton, C. Thaung, J.A. Stevenson, R. Arkell, P. Mburu, R. Hardisty, A. Kiernan, A. Erven, K.P. Steel, S. Voegeling, J.L. Guenet, C. Nickols, R. Sadri, M. Nasse, A. Isaacs, K. Davies, M. Browne, E.M. Fisher, J. Martin, S. Rastan, S.D. Brown, and J. Hunter. 2000. A systematic, genome-wide, phenotype-driven mutagenesis programme for gene function studies in the mouse. Nat Genet. 25:440–443.

Oleynikov, Y., and R.H. Singer. 2003. Real-time visualization of ZBP1 association with beta-actin mRNA during transcription and localization. Curr Biol. 13:199–207.

Oshima, K., M. Takeda, E. Kuranaga, R. Ueda, T. Aigaki, M. Miura, and S. Hayashi. 2006. IKK epsilon regulates F actin assembly and interacts with Drosophila IAP1 in cellular morphogenesis. Curr Biol. 16:1531–1537.

Palacios, I.M. 2007. How does an mRNA find its way? Intracellular localisation of transcripts. Semin Cell Dev Biol. 18:163–170.

Palazzo, R.E., J.M. Vogel, B.J. Schnackenberg, D.R. Hull, and X. Wu. 2000. Centrosome maturation. Curr Top Dev Biol. 49:449–470.

Pandey, R., S. Heeger, and C.F. Lehner. 2007. Rapid effects of acute anoxia on spindle kinetochore interactions activate the mitotic spindle checkpoint. J Cell Sci. 120:2807–2818.

Port, F., and S.L. Bullock. 2016. Creating Heritable Mutations in Drosophila with CRISPR-Cas9. Methods Mol Biol. 1478:145–160.

Port, F., H.M. Chen, T. Lee, and S.L. Bullock. 2014. Optimized CRISPR/Cas tools for efficient germline and somatic genome engineering in Drosophila. Proc Natl Acad Sci U S A. 111:E2967–2976.

Raff, J.W., W.G. Whitfield, and D.M. Glover. 1990. Two distinct mechanisms localise cyclin B transcripts in syncytial Drosophila embryos. Development. 110:1249–1261.

Reck-Peterson, S.L., W.B. Redwine, R.D. Vale, and A.P. Carter. 2018. The cytoplasmic dynein transport machinery and its many cargoes. Nat Rev Mol Cell Biol. 19:382–398.

Robert, X., and P. Gouet. 2014. Deciphering key features in protein structures with the new ENDscript server. Nucleic Acids Res. 42:W320–324.

Ryder, P.V., J. Fang, and D.A. Lerit. 2020. centrocortin RNA localization to centrosomes is regulated by FMRP and facilitates error-free mitosis. J Cell Biol. 219.

Ryder, P.V., and D.A. Lerit. 2018. RNA localization regulates diverse and dynamic cellular processes. Traffic. 19:496–502.

Ryder, P.V., and D.A. Lerit. 2020. Quantitative analysis of subcellular distributions with an open-source, object-based tool. Biol Open. 9.

Safieddine, A., E. Coleno, S. Salloum, A. Imbert, A.M. Traboulsi, O.S. Kwon, F. Lionneton, V. Georget, M.C. Robert, T. Gostan, C.H. Lecellier, R. Chouaib, X. Pichon, H. Le Hir, K. Zibara, F. Mueller, T. Walter, M. Peter, and E. Bertrand. 2021. A choreography of centrosomal mRNAs reveals a conserved localization mechanism involving active polysome transport. Nat Commun. 12:1352.

Salvador-Garcia, D., L. Jin, A. Hensley, M. Golcuk, E. Gallaud, S. Chaaban, F. Port, A. Vagnoni, V.J. Planelles-Herrero, M.A. McClintock, E. Derivery, A.P. Carter, R. Giet, M. Gur, A. Yildiz, and S.L. Bullock. 2023. A force-sensitive mutation reveals a spindle assembly checkpoint-independent role for dynein in anaphase progression. bioRxiv.

Schindelin, J., I. Arganda-Carreras, E. Frise, V. Kaynig, M. Longair, T. Pietzsch, S. Preibisch, C. Rueden, S. Saalfeld, B. Schmid, J.Y. Tinevez, D.J. White, V. Hartenstein, K. Eliceiri, P. Tomancak, and A. Cardona. 2012. Fiji: an open-source platform for biological-image analysis. Nat Methods. 9:676-682.

Schlager, M.A., A. Serra-Marques, I. Grigoriev, L.F. Gumy, M. Esteves da Silva, P.S. Wulf, A. Akhmanova, and C.C. Hoogenraad. 2014. Bicaudal d family adaptor proteins control the velocity of Dynein-based movements. Cell Rep. 8:1248–1256.

Schneider-Poetsch, T., J. Ju, D.E. Eyler, Y. Dang, S. Bhat, W.C. Merrick, R. Green, B. Shen, and J.O. Liu. 2010. Inhibition of eukaryotic translation elongation by cycloheximide and lactimidomycin. Nat Chem Biol. 6:209–217.

Sepulveda, G., M. Antkowiak, I. Brust-Mascher, K. Mahe, T. Ou, N.M. Castro, L.N. Christensen, L. Cheung, X. Jiang, D. Yoon, B. Huang, and L.E. Jao. 2018. Co-translational protein targeting facilitates centrosomal recruitment of PCNT during centrosome maturation in vertebrates. Elife. 7.

Sladewski, T.E., N. Billington, M.Y. Ali, C.S. Bookwalter, H. Lu, E.B. Krementsova, T.A. Schroer, and K.M. Trybus. 2018. Recruitment of two dyneins to an mRNA-dependent Bicaudal D transport complex. Elife. 7.

Soltys, B.J., and G.G. Borisy. 1985. Polymerization of tubulin in vivo: direct evidence for assembly onto microtubule ends and from centrosomes. J Cell Biol. 100:1682–1689.

Suter, B., and R. Steward. 1991. Requirement for phosphorylation and localization of the Bicaudal-D protein in Drosophila oocyte differentiation. Cell. 67:917–926.

Tadros, W., and H.D. Lipshitz. 2009. The maternal-to-zygotic transition: a play in two acts. Development. 136:3033–3042.

Theurkauf, W.E. 1994. Immunofluorescence analysis of the cytoskeleton during oogenesis and early embryogenesis. Methods Cell Biol. 44:489–505.

UniProt, C. 2023. UniProt: the Universal Protein Knowledgebase in 2023. Nucleic Acids Res. 51:D523–D531.

Veeranan-Karmegam, R., D.P. Boggupalli, G. Liu, and G.B. Gonsalvez. 2016. A new isoform of Drosophila non-muscle Tropomyosin 1 interacts with Kinesin-1 and functions in oskar mRNA localization. J Cell Sci. 129:4252–4264.

Vertii, A., H. Hehnly, and S. Doxsey. 2016. The Centrosome, a Multitalented Renaissance Organelle. Cold Spring Harb Perspect Biol. 8.

Zein-Sabatto, H., and D.A. Lerit. 2021. The Identification and Functional Analysis of mRNA Localizing to Centrosomes. Front Cell Dev Biol. 9:782802.

